# Synthetic genomic dissection of enhancer context sensitivity and synergy

**DOI:** 10.1101/2025.08.13.669251

**Authors:** Raquel Ordoñez, André M. Ribeiro-dos-Santos, Colleen McLoughlin, Gwen Ellis, Hannah J. Ashe, Nerea Berastegui Zufiaurre, Ran Brosh, Milosz Majewski, Kelly Barriball, Brendan Camellato, Jef D. Boeke, Matthew T. Maurano

## Abstract

Noncoding disease and trait-associated genetic variation is frequently interpreted in the context of genomic regulatory elements such as DNase I hypersensitive sites (DHSs). But while most DHSs lie within a few kilobases of another DHS, regulatory elements are typically analyzed individually without accounting for their neighbors. We characterize multiple heterotypic DHS combinations from different critical mESC regulator loci, all delivered in a constant chromosomal context replacing the *Sox2* Locus Control Region (LCR). We employ an optimized high-throughput multiplexed delivery pipeline enabling analysis of 213 distinct payloads in 641 mouse embryonic stem cell (mESC) clones. We identify widespread examples of context-dependent enhancers which have no activity on their own but can more than double the activity of a neighboring DHS. Enhancers exhibit synergy only with certain partners, and deliveries to the *Igf2*/*H19* locus show that synergy is not constrained to a single genomic context. We further show that synergy between neighboring DHSs decays as a characteristic function of distance, with its influence extending up to 4 kilobases. We fine map this context dependency to the contribution of individual transcription factor recognition sequences. Our approach implicates the specific sequence and architectural features underpinning pervasive genomic context effects, and outlines a direction for modeling the functional impact of noncoding regulatory variation on common human traits and diseases.

## INTRODUCTION

Gene expression is tightly regulated by a complex network of enhancers which interact with gene promoters over long genomic distances. Enhancers were originally defined as short modular DNA sequences capable of autonomously driving promoter transcription regardless of their distance or orientation (Banerji et al. 1981; Schaffner 2015). In the genomic age, the term enhancer has come to include genomic sites identified by the presence of biochemical hallmarks including accessibility to DNase I or other enzymes, characteristic histone modifications, and enhancer RNA (eRNA) transcription (Catarino and Stark 2018). Yet there is an incomplete overlap between these definitions, and only a fraction of the such genomically identified elements show enhancer activity in reporter assays (The ENCODE Project Consortium et al. 2020). A key open question is how their surrounding context modulates enhancer function. While enhancers are usually considered as individual sequences, multiple genomic regulatory elements are frequently found in close proximity, including at super-enhancers (Whyte et al. 2013) or locus control regions (LCR) (Li et al. 2002). Moreover, genome-wide association studies have shown that noncoding disease-associated variants are preferentially enriched within regulatory DNA (Maurano et al. 2012), with particular enrichment in densely clustered enhancer regions (Parker et al. 2013; Hnisz et al. 2013), underscoring the importance of understanding these regulatory landscapes.

Understanding how the regulatory logic of an enhancer cluster is encoded by the underlying DNA sequence requires a perturbation approach that can test how different combinations of regulatory elements interact in a chromosomal context. Synthetic regulatory genomics addresses this challenge by enabling perturbational studies of complex regulatory loci (Maurano 2026). For example, Big-IN employs recombinase-mediated integration at a pre-engineered landing pad to facilitate scarless genomic insertion of engineered payloads sized on the order of 100 kilobases (Brosh et al. 2021). This provides a powerful framework to systematically reconstruct and perturb entire regulatory landscapes, making it possible to test how combinations of regulatory elements encode transcriptional outcomes.

Synthetic regulatory genomics has been used to decompose the regulatory architecture of the *Sox2* locus, which encodes a key regulator of pluripotency, reprogramming, early development, and that is also a major driver in many cancer types. *Sox2* expression in mouse embryonic stem cells (mESC) requires a distal LCR (Zhou et al. 2014; Li et al. 2014). Dissection of the *Sox2* LCR has revealed extensive context-dependence among its constituent DNase I hypersensitive sites (DHSs) (Brosh et al. 2023). For example, *Sox2* DHS24 acts as an autonomous enhancer while its neighbor DHS23 has no detectable activity alone but can double the activity of DHS24 when the two are placed nearby (Brosh et al. 2023). Other studies addressing sufficiency in a native genomic context have identified genomic elements without autonomous activity that synergize with nearby enhancers (Thomas et al. 2021; Brosh et al. 2023; Blayney et al. 2023) or sequences that contribute to enhancer-promoter communication through architectural interactions (Batut et al. 2022; Chen et al. 2023; Bower et al. 2025). Regulatory elements can interact redundantly, additively, or synergistically, so that their combined activity exceed the sum of their individual effects (Bothma et al. 2015; Thomas et al. 2021; Choi et al. 2021; Martinez-Ara et al. 2022; Lin et al. 2022; Brosh et al. 2023; Blayney et al. 2023; Zhou et al. 2026; Koeppel et al. 2025). Together, this suggests that enhancers may be composite regulatory units whose function depends on context and cooperation.

Here we systematically investigate enhancer context dependence using a synthetic regulatory genomics approach. We investigate DHS combinations from *Sox2* and other key mESC regulatory loci and profile their ability to activate gene expression at long range. We identify widespread context dependence, characterize the impact of genomic distance on enhancer interaction, and fine-map the sequence determinants of synergy to specific transcription factor (TF) motif arrangements. Our findings suggest that regulatory DNA function must be seen through the lens of context effects from neighboring regulatory elements, and implicate specific sequence and architectural context for incorporation into models of regulatory variation in human traits and disease.

## RESULTS

### A multiplexed delivery platform for synthetic enhancer analysis

To systematically dissect enhancer context dependence, we developed a scalable system capable of evaluating a high number of synthetic enhancer constructs in a controlled sequence environment. We observed that Big-IN delivery of payloads <20 kb benefit from high transfection efficiency and rarely demonstrate internal rearrangements or deletions, regularly yielding large numbers of colonies with high genotyping and Capture-seq success rates. We therefore designed a barcoding experiment to assess whether Big-IN can support multiplexed delivery of short payloads. We built a 2.4-kb payload library including a barcode to uniquely identify independent insertions and a GFP expression cassette. We delivered this library to landing pads at *Igf2/H19* and *Hprt* loci, counter-selected for successful delivery, and quantified barcode diversity specifically from on-target deliveries using an amplicon-based deep sequencing assay (**Fig. S1a**). Sequencing revealed hundreds of uniquely barcoded on-target payload integrations both at the *Igf2/H19* and *Hprt* loci (**Fig. S1b**). Fluorescent imaging showed that >90% of colonies showed GFP signal, confirming the efficiency of payload delivery (**Fig. S1c**). These results demonstrate that Big-IN can be used for multiplexed delivery of hundreds of different assemblies.

Encouraged by the feasibility of pooled delivery, we sought to establish a scalable workflow for multiplexed Big-IN experiments (**Fig. 1**). First, to enable efficient cloning of multiple Big-IN payloads, we used Golden Gate assembly for the construction of a diverse library of payloads (**Table S1**, **Table S2**, **Data S1**). We developed a new pINEAPLE (plasmid for INnovativE Assembly of PayLoads in *E. coli*) vector specifically to facilitate scalable assembly and delivery of Big-IN payloads. Compared to the original pLM1110 backbone designed for yeast assembly of large payloads (Brosh et al. 2021), it is significantly smaller (5.5 kb vs. 14 kb), supports high-copy growth in standard bacteria strains, incorporates a green/white GFP-based screening system, includes a simplified mammalian selection cassette, and has modular design features to allow swapping between both pLM1110 and pINEAPLE vector systems. Each payload was flanked by a pair of unique nucleotide sequences (UNS) to facilitate genotyping after delivery. This system enabled consistent and scalable preparation of multiplexed Big-IN payloads suitable for pooled delivery.

**Fig. 1.**
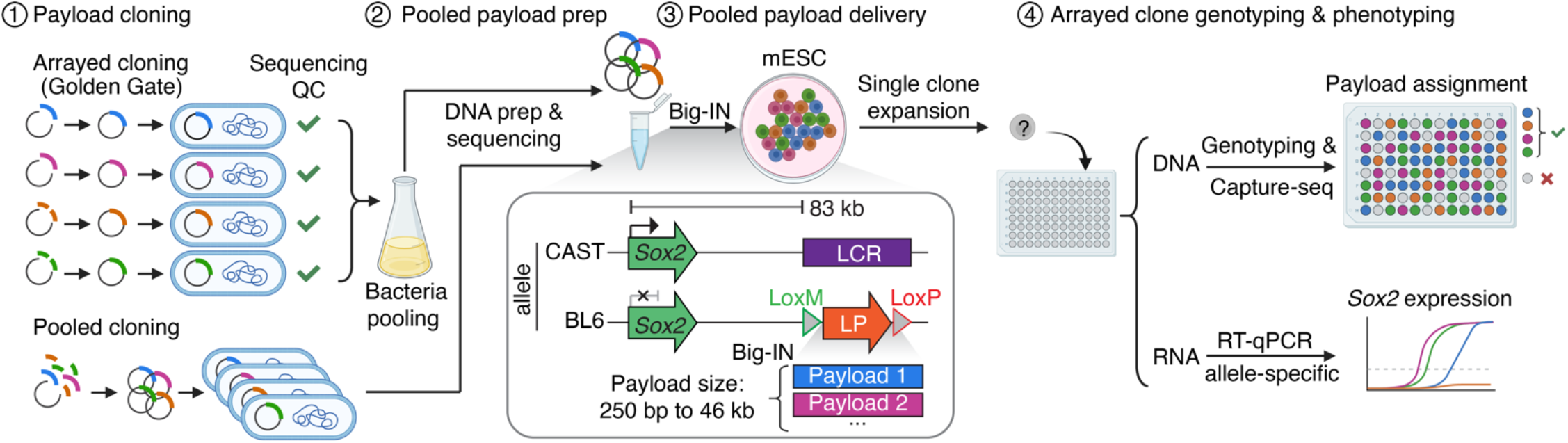
Multiplexed Big-IN approach. Schematic of pooled-to-arrayed strategy for scalable Big-IN payload delivery and phenotyping. Payloads are assembled either by arrayed Golden Gate Assembly followed by pooling, or direct pooled cloning. Pooled payload DNA is then extracted and delivered to mESC using Big-IN. At the *Sox2* locus, payloads are delivered to a landing pad (LP) installed in place of the *Sox2* Locus Control Region (LCR), such that *Sox2* expression from the engineered allele is driven by the delivered payload. The distance between the delivered payloads and the *Sox2* promoter is indicated. Following selection on pools, individual clones are expanded in arrayed 96-well format. PCR genotyping and Capture-seq are used to assign payload identity. Allele-specific qRT-PCR is performed to quantify expression from the engineered allele in each clone. Some icons are from BioRender.

We then developed a scalable readout strategy using plate-based phenotyping (**Fig. 1**). We delivered payloads to LP-LCR C57BL6/6J x CAST/EiJ (BL6xCAST) F1 hybrid mESCs containing a Big-IN landing pad replacing the *Sox2* LCR on their BL6 allele (Brosh et al. 2023). In this system, *Sox2* expression from the engineered allele depends on long-range activation from the delivered payloads. Pooled Big-IN payload libraries were delivered in a single transfection (**Table S3**), followed by selection for correctly delivered clones in pooled format. Culturing cells in pooled format through selection significantly reduced cell culture requirements. Clones were then picked and grown in a 96-well plate (**Table S4**), genotyped for on-target integration using PCR, and then profiled for expression using RT-qPCR (**Table S5**, **Table S6, Data S2**). Expression from the engineered allele (BL6) was quantified relative to the unedited CAST allele as an internal control. To confirm correct single-copy on-target delivery and identify the payload delivered in each clone, we employed plate-based genomic DNA extraction and library construction, followed by targeted capture sequencing (Brosh et al. 2021). We developed a payload deconvolution strategy that combines three orthogonal analysis approaches: junction mapping using a modified version of *bamintersect* (Brosh et al. 2021), read coverage analysis, and unique sequence detection (k-mer genotyping) (**Table 1** and **Table S7**). This enabled flexible deconvolution across different payload library designs. Approximately 60-90% of delivery clones isolated passed PCR genotyping, confirming successful on-target integration. Of these, 80-90% passed subsequent Capture-seq validation to exclude off-target or incomplete integration, payload backbone or pCAGiCre integration, or landing pad retention, and were successfully genotyped for payload identity. Together, this framework establishes a scalable platform for generation, pooled delivery, validation, deconvolution, and phenotyping of synthetic sequences.

**Table 1.**
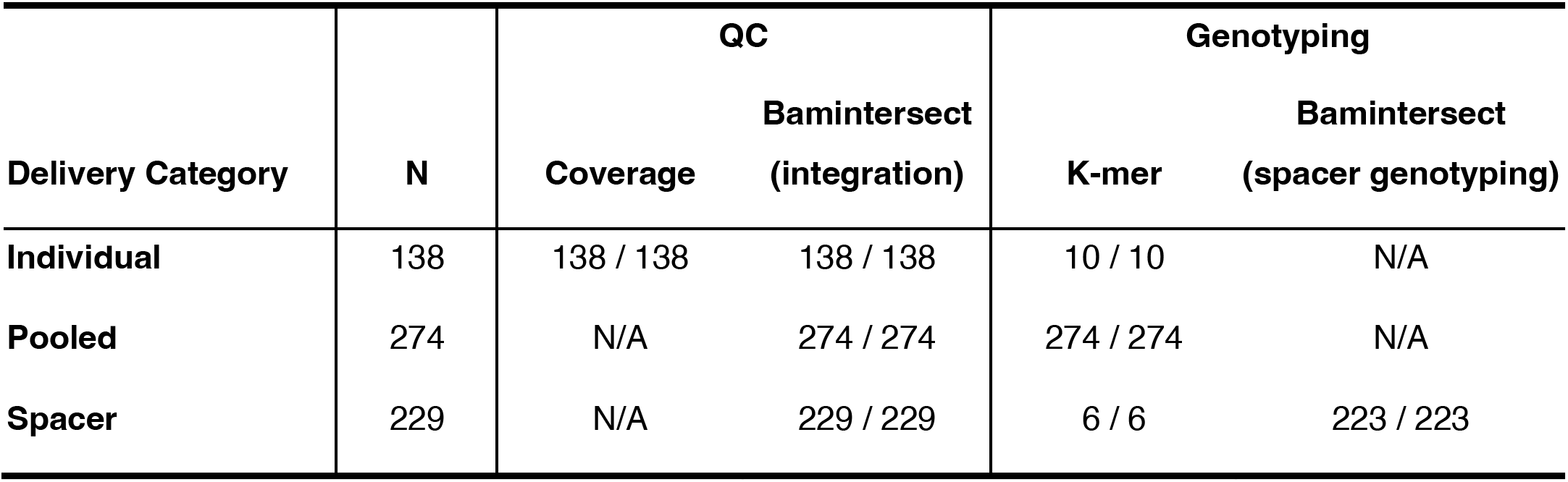
Engineered mESC clone quality control and genotyping summary. mESC clones were verified by analysis of sequencing coverage depth, and sequence composition according to their category (Individual, Pooled, or Spacer delivery). Individually delivered clones were verified by read coverage at defined regions. All payload integrations were verified by bamintersect. Clones from pooled deliveries, individually delivered spacer clones, and clones inferred to have small spacers were verified by k-mer genotyping. Spacer payload delivery was verified by bamintersect spacer genotyping. See details in **Methods**. Under each analysis are indicated the number of successful clones and the number of clones verified. N/A, not applicable.

| Delivery Category | N | QC |  | Genotyping |  |
| --- | --- | --- | --- | --- | --- |
|  |  | Coverage | Bamintersect<br>(integration) | K-mer | Bamintersect<br>(spacer genotyping) |
| Individual | 138 | 138 / 138 | 138 / 138 | 10 / 10 | N/A |
| Pooled | 274 | N/A | 274 / 274 | 274 / 274 | N/A |
| Spacer | 229 | N/A | 229 / 229 | 6 / 6 | 223 / 223 |

### Combinatorial DHS libraries show widespread enhancer cluster synergy

We applied this platform to test whether context-dependent activity is a general feature of developmental enhancers. We selected candidate DHSs from the *Nanog*, *Sall1*, and *Prdm14* loci, each representing key mESC regulator loci previously characterized through deletion analysis (Hnisz et al. 2015; Blinka et al. 2016; Moorthy et al. 2017; Singh et al. 2021; Vos et al. 2021) (**Fig. 2a-d**). We cloned candidate DHSs individually or in pairs with *Sox2* 23 (previously classified as con-text-dependent) and *Sox2* 24 (autonomous), synthetic sequences derived from *Sox2* DHS23 and *Sox2* DHS24 that omit or shorten a small number of repetitive regions to facilitate DNA synthesis (Brosh et al. 2023). Candidate payloads were delivered in place of the *Sox2* LCR and verified mESC clones were profiled for *Sox2* expression (**Fig. 2e, Fig. S2**). None of the candidates had strong activity on their own, and all were weaker than *Sox2* 24 (**Fig. S3a**). But some DHS pairs had significantly higher activity than the constituent DHSs alone.

**Fig. 2.**
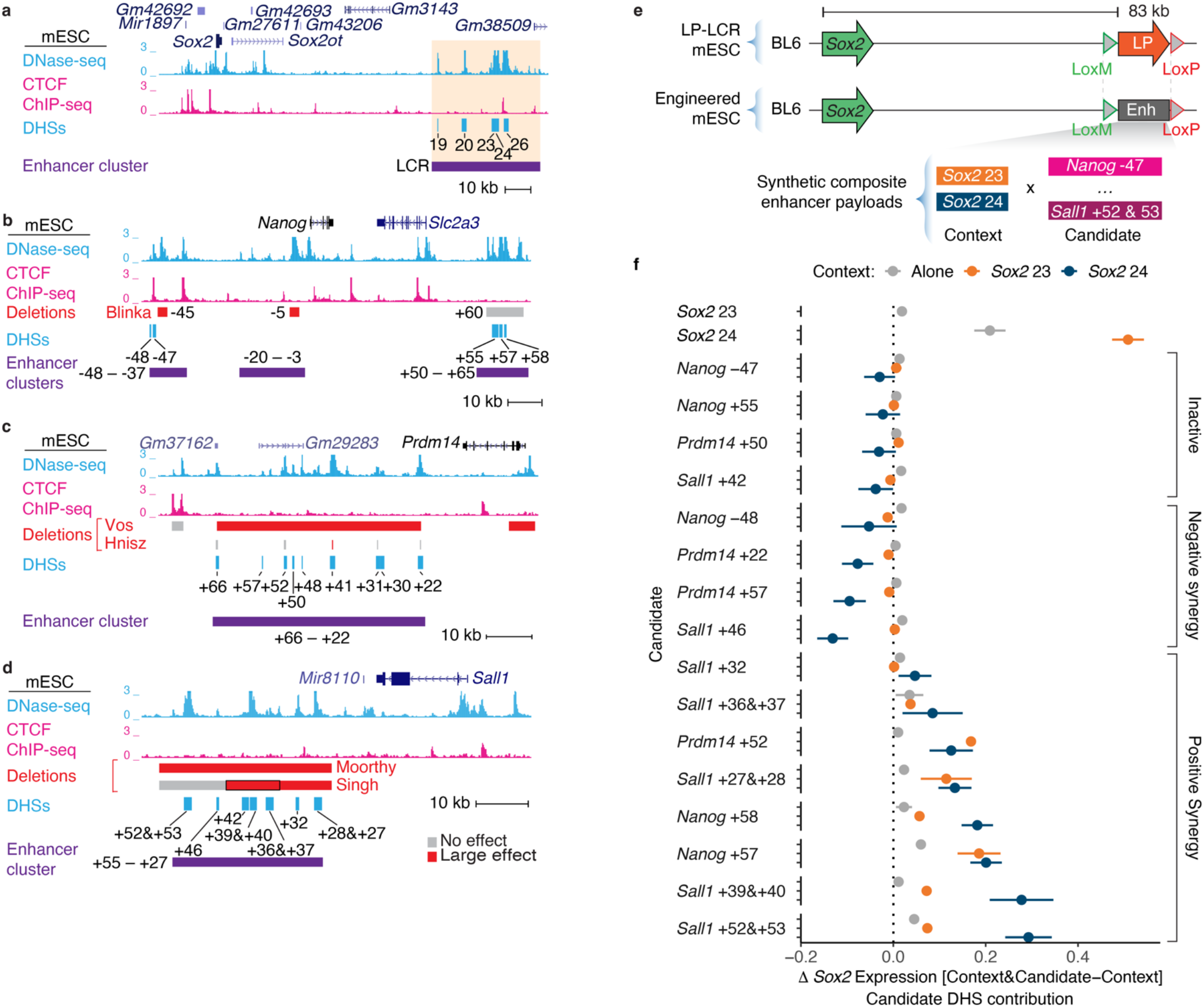
**Identification of novel context-sensitive DHSs. a-d**. *Sox2* (**a**), *Nanog* (**b**), *Prdm14* (**c**), and *Sall1* (**d**) loci showing mESC DNase-seq and CTCF ChIP-seq. Red indicates deletions reported to affect expression of the target gene, gray indicates no effect (Hnisz et al. 2015; Blinka et al. 2016; Moorthy et al. 2017; Singh et al. 2021; Vos et al. 2021). Enhancer cluster boundaries are marked with purple boxes. Orange shading in (**a**) denotes the Big-IN landing pad replacing the BL6 allele of the *Sox2* LCR. **e**. Schematic of strategy to identify enhancer context sensitivity delivering payloads to replace the *Sox2* LCR. **f.** Expression analysis of candidate DHSs tested alone (gray), or in combination with *Sox2* 23 (orange) or *Sox2* 24 (blue). Contribution of each DHS, computed as the difference in *Sox2* expression between payload pairs differing solely by the presence of the candidate DHS. Points indicate mean expression change. Error bars indicate SD across all pairwise combinations of clones.

We explored how each individual context DHS (*Sox2* 23 or *Sox2* 24) modulated enhancer activity by comparing each of the candidate DHSs to the activity of the payload without the candidate (**Fig. 2f, Fig. S3b**). This analysis resolved multiple categories of enhancer context dependence: (i) DHSs that show no activity at all, whether on their own or with a partner DHS; (ii) DHSs that show no activity on their own but synergize negatively with *Sox2* 24; (iii) DHSs that synergize specifically with *Sox2* 24; or (iv) with both *Sox2* 23 and *Sox2* 24. Thus, the differences observed among candidate DHSs when paired with *Sox2* 23 or *Sox2* 24 indicate that the regulatory contribution of a given DHS depends on its partner.

To identify genomic predictors of DHS synergy, we examined different biochemical hallmarks of regulatory activity in the candidate DHSs, such as TF occupancy patterns, histone modifications, PRO-seq signal, STARR-seq reporter assay activity, and sequence conservation (**Fig. S4**). None of these features alone reliably predicted the presence or direction of synergy. Notably, some DHSs with high STARR-seq activity failed to behave as autonomous enhancers in our genomic context, such as *Sall1* +39&+40 and *Sall1* +27&+28, highlighting that reporter-based activity does not necessarily translate to autonomous function in a different context. We next asked whether specific TF motif families were associated with the presence or direction of synergy. TF motif families showed broadly similar distributions across all candidate DHSs, with no clear enrichment of any specific family in a given category (**Fig. S5**), suggesting that synergy is not driven by a single dominant TF motif family.

### DHS synergy operates outside the *Sox2* locus

We next tested whether DHS synergy could be detected outside the *Sox2* locus. The imprinted *Igf2/H19* locus is the paradigmatic example of a regulatory switch: differential CTCF occupancy at the imprinting control region (ICR) directs the target of a distal enhancer cluster to either *H19* or *Igf2* (Bell and Felsenfeld 2000; Hark et al. 2000; Ordoñez et al. 2024). While the native *H19* enhancer cluster is inactive in mESC, the *Sox2* LCR can activate both genes in mESC (Ordoñez et al. 2024). We generated payloads ranging in size from 116 to 143 kb to rewrite the *Igf2/H19* locus and replace the *H19* enhancer cluster with DHSs from the *Sox2* and the *Sall1* loci. All payloads included deletion of all 6 CTCF binding sites within the imprinting control region (ICR) (ΔCTCF-ICR) which enables the *H19* enhancers to activate the more distal *Igf2* gene (Ordoñez et al. 2024). Payloads were delivered to LP-*Igf2*/*H19* mESC harboring a Big-IN landing pad replacing the BL6 allele of the *Igf2*/*H19* locus (**Fig. 3a-b**). mESC clones were subsequently verified and profiled for expression of both *Igf2* and *H19* from the engineered allele using allele-specific RT-qPCR.

**Fig. 3.**
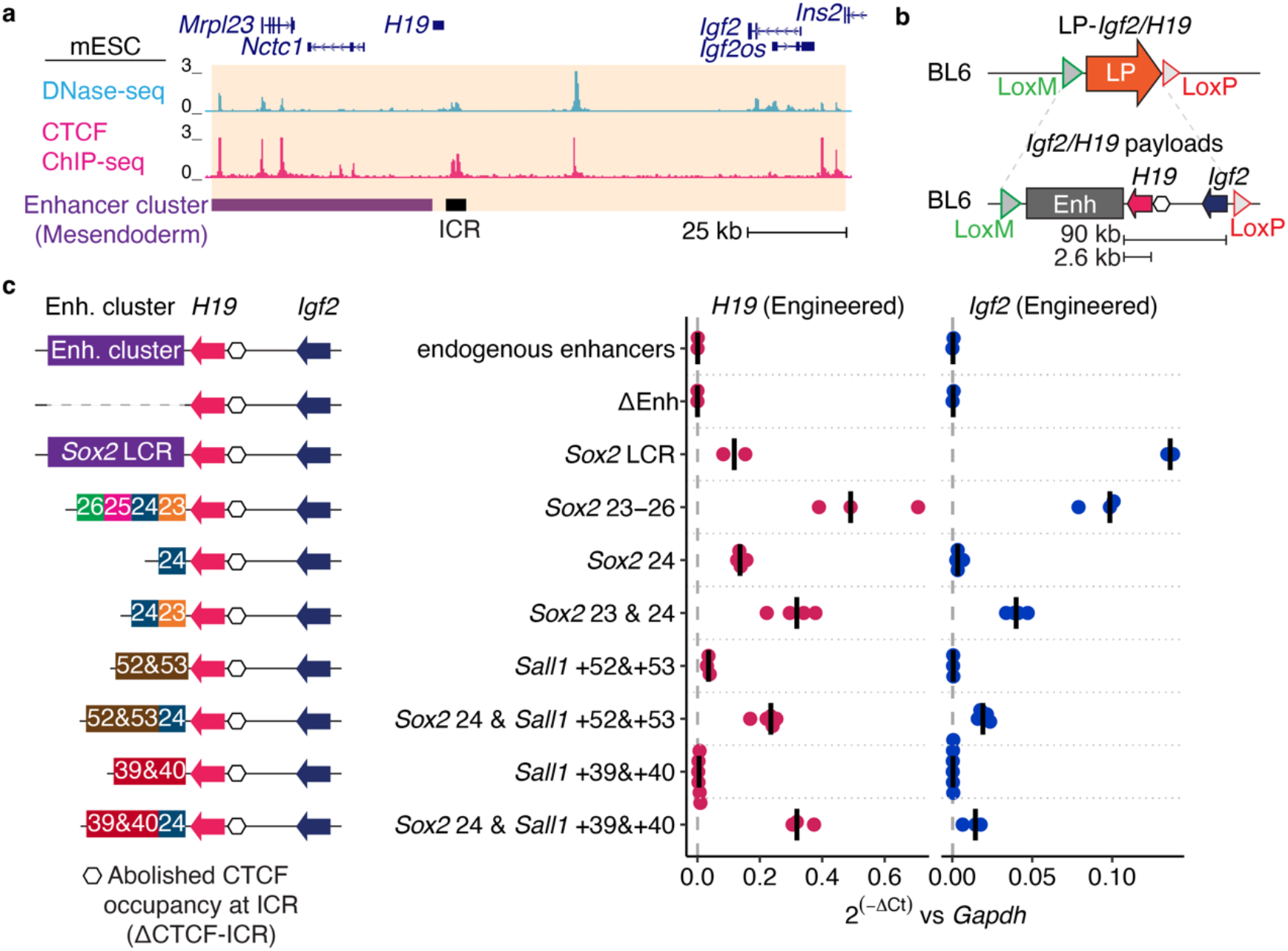
Enhancer DHS synergy is preserved at the *Igf2/H19* locus. **a.** *Igf2* locus showing mESC DNase-seq and CTCF ChIP-seq. The Big-IN landing pad replacing the BL6 allele of the *Igf2/H19* locus is denoted by orange shading. The *H19* enhancer cluster replaced by candidate sequences is marked with a purple box, and the imprinting control region (ICR) with a black box. **b**. Schematic of strategy to deliver payloads to replace the whole *Igf2/H19* locus in the BL6 allele with assemblies including the synthetic enhancer clusters. The distance between the delivered payloads and the *H19* and *Igf2* promoters is indicated. All payloads included deletion of all 6 CTCF binding sites within the ICR (empty hexagon, ΔCTCF-ICR), abolishing CTCF occupancy and allowing the enhancer cluster to interact with both gene promoters (Ordoñez et al. 2024). **c**. *Igf2*/*H19* expression analysis for enhancer clusters delivered replacing the endogenous enhancer cluster. Each point represents the expression from the engineered allele in an independent mESC clone. Bars indicate median. Expression from the engineered allele was normalized to *Gapdh*. Vertical dashed line indicates median expression of ΔEnh. Labels indicate DHS numbers as in Fig. 2a**-d**. The *Sox2* LCR payload includes the full-length enhancer sequence (Brosh et al. 2023). Clones for endogenous enhancers, ΔEnh and *Sox2* LCR were previously reported (Ordoñez et al. 2024).

As previously shown, the full *Sox2* LCR activated both *Igf2* and *H19*. Compared to the full *Sox2* LCR (41 kb), the core LCR (a 6.3-kb region including *Sox2* 23–26) drove lower *Igf2* expression but higher *H19* expression (**Fig. 3c**). *Sox2* 24 showed even lower activation of *H19*, and none of *Igf2*. But *Sox2* 23 & 24 showed higher activation of both genes, behaving more like *Sox2* 23–26. Thus, DHS synergy within the *Sox2* LCR operates beyond its endogenous locus, and different patterns of synergy can redirect its target genes.

Examining the candidate DHSs from the *Sall1* locus, neither *Sall1* +52&+53 or *Sall1* +39&+40 showed any activity on their own, but each behaved similar to *Sox2* 23 in that they synergized with *Sox2* 24 to activate both *H19* and *Igf2*. These results reinforce that DHS synergy is not private to the *Sox2* locus (**Fig. 3c**).

### Full enhancer clusters show lower activity than closely linked DHSs

Given the large differences between the *Sox2* LCR and its constituent DHSs to activate *Igf2* and *H19*, we investigated more deeply the functional differences between other complex enhancer clusters and their constituent regulatory elements. We engineered payloads ranging from 6 to 45 kb in size encompassing intact enhancer clusters and including multiple DHSs and the intervening sequences and delivered them replacing the *Sox2* LCR. All enhancer clusters except *Nanog* -48 – -37 drove *Sox2* expression, but at very modest levels: all were below the contribution of the single DHS *Sox2* 24 (**Fig. 4**). The *Nanog* -48 – -37 cluster was inactive both as a unit and as individual elements (*Nanog* -48 and *Nanog* -47, **Fig. 2f**). *Prdm14* +66 – +22 and *Sall1* +55 – +27 drove expression of their endogenous target genes at 20% and 34% of native *Sox2* expression respectively (**Fig. S6**), suggesting that their activity at the *Sox2* locus was lower than at their native loci. Surprisingly given the limited activity of the *Nanog* +50 – +65 enhancer cluster, *Nanog* is expressed 3.7 times higher than *Sox2*, and *Nanog* +57 and *Nanog* +58 showed strong synergy when paired individually with *Sox2* 23 or *Sox2* 24 (**Fig. 2f**).

**Fig. 4.**
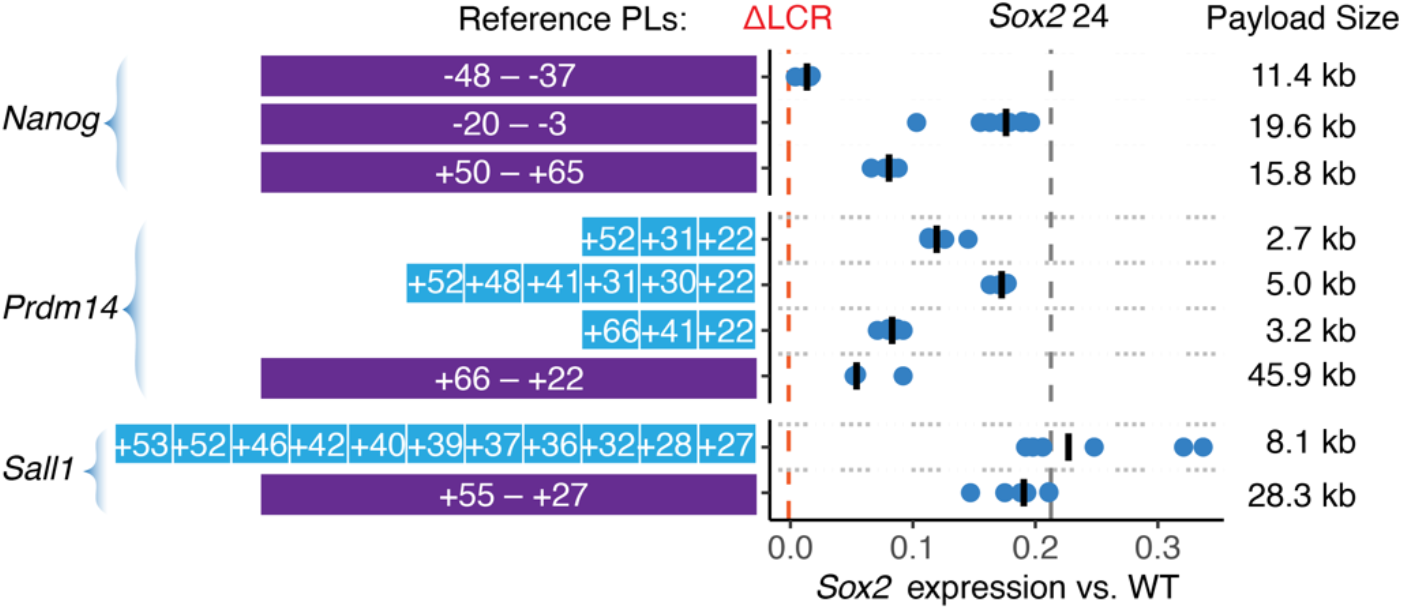
*Sox2* expression analysis for enhancer clusters delivered replacing the *Sox2* LCR. Payloads include full-length enhancer sequences (including native intervening sequences, represented as purple blocks) and collapsed enhancer clusters (DHSs assembled together without the intervening sequences, represented with blue blocks). Each point represents *Sox2* expression from the engineered allele in an independent mESC clone. Bars indicate median. Expression from the BL6 allele was normalized to the CAST allele and scaled between 0 (Δ*Sox2*) to 1 (WT LCR). Vertical dashed lines indicate median expression of baseline payloads. Labels indicate DHS numbers as in Fig. 2a**-d**.

Given the relatively low activity of these intact enhancer clusters, we investigated “collapsed” enhancer clusters which included multiple DHSs linked in close proximity without intervening sequence. Despite comprising only a subset of the DHSs present in the full sequences, some collapsed enhancer clusters drove higher *Sox2* expression than the full enhancer clusters (**Fig. 4**). One possible explanation is the full enhancer clusters include elements conferring both positive and negative activity, and overall activity reflects the net outcome of these effects. For example, both *Prdm14* +57 and *Prdm14* +22 functioned as repressors with *Sox2* 24. However, another possibility is that extended spacing between DHSs diminishes their interaction.

### DHS pair spacing modulates enhancer synergy

Given these results suggesting that reduced DHS spacing could increase enhancer cluster activity, we sought to directly isolate the effect of spacing on DHS pair activity. To dissect DHS spacing independently of the function of the intervening sequence, we employed SynHPRT1R, a synthetic payload representing a “backwards” version (i.e., reversed but not complemented) of the 101-kb human *HPRT1* locus (Camellato et al. 2024). This DNA preserves certain features of the native *HPRT1* locus, such as base composition and repetitive sequence, while disrupting higher-order features such as coding sequence and TF recognition sites (except palindromes). SynHPRT1R has previously been demonstrated to have no ATAC-seq or H3K27ac signal upon delivery into mESC (Camellato et al. 2024) (**Fig. S7a**). We employed a pooled library cloning strategy using a ladder of BsiWI and AvrII-digested SynHPRT1R fragments ranging from 247 bp to 4065 bp (**Fig. 5a**, **Fig. S7a**). Sourcing the library from restriction-enzyme digests ensures that fragments with closely related sizes have non-overlapping and independent sequences (**Fig. S7a**). Upon delivery replacing the *Sox2* LCR, Capture-seq was used to deconvolute the spacer identity for each mESC clone.

**Fig. 5.**
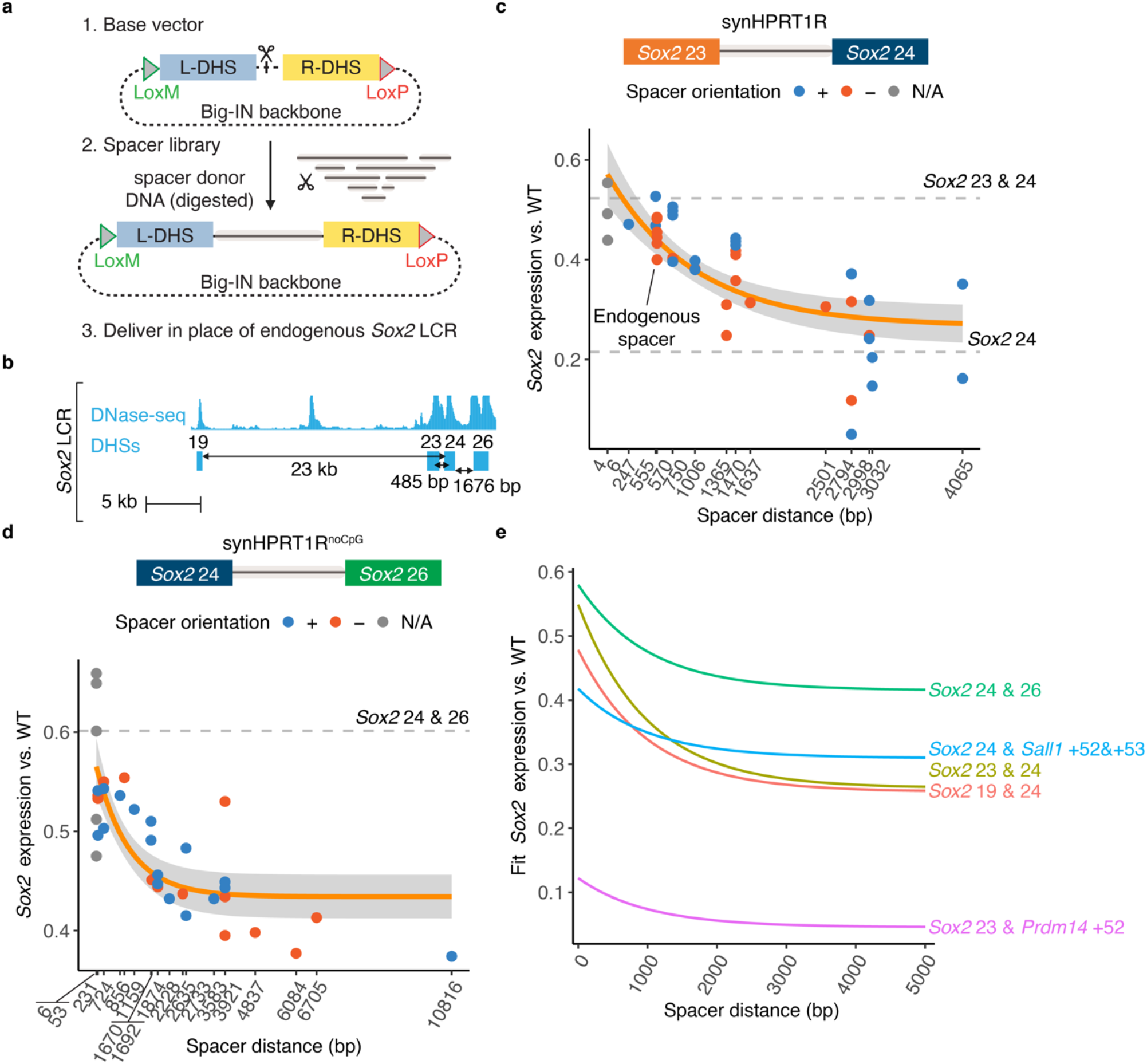
Synergistic enhancer function depends on distance between DHSs. **a**. Schematic of spacing library cloning strategy: spacer fragments (6 bp–10 kb) derived from restriction-digested donor BAC DNA were cloned between a pair of DHSs and integrated using Big-IN to replace the endogenous *Sox2* LCR. Individual clones were isolated for analysis. **b**. Schematic of the *Sox2* LCR highlighting DHSs included in spacer experiments. Distances represent endogenous spacing between DHSs. **c-d**. *Sox2* expression analysis using qRT-PCR including DHS pairs *Sox2* 23 and *Sox2* 24 (**c**) and *Sox2* 24 and *Sox2* 26 (**d**), separated by spacers from digests of SynHPRT1R (**c**) and SynHPRT1R^noCpG^ (**d**). Each point represents *Sox2* expression from the engineered allele in an independent mESC clone. Expression from the BL6 allele was normalized to the CAST allele and scaled between 0 (Δ*Sox2*) to 1 (WT LCR). Orange line indicates exponential fit with 95% confidence interval shaded in gray. Horizontal dashed lines indicate median expression for baseline payloads. Color represents spacer orientation; N/A (grey) indicates that no spacer was cloned between the two DHSs and only the restriction enzyme sequence remains. **e**. Comparison of distance dependence for multiple DHS pairs from **c-d** and **Fig. S9**. Lines represent exponential fits.

We first assessed a library of SynHPRT1R spacer fragments cloned between *Sox2* 23 and *Sox2* 24. The results showed that the synergy between *Sox2* 23 and *Sox2* 24 decays with increasing distance. *Sox2* expression decreased monotonically to the activity of *Sox2* 24 alone as the spacer distance increased to 4 kb (**Fig. 5b-c**), consistent with a functional dissociation of both DHSs.

To investigate whether this distance-dependent synergy also affected autonomous DHSs, we investigated *Sox2* 24 and *Sox2* 26, both of which show autonomous activation of *Sox2* (Brosh et al. 2023). To further generalize our spacer approach, we tested our strategy with SynHPRT1R^noCpG^, a derivative of SynHPRT1R wherein CpGs were mutated away and which lacks ATAC-seq, H3K27ac signal, and H3K27me3 signal in mESC (Camellato et al. 2024) (**Fig. S7b**). AvrII-digested synHRPT1R^noCpG^ fragments ranging from 53 bp to 10.8 kb were cloned between *Sox2* 24 and *Sox2* 26, a DHS pair spaced 1.7 kb apart in the genome that also exhibits synergy when placed more closely. These data showed a similar distance dependence between *Sox2* 24 and *Sox2* 26 (**Fig. 5d**). These results demonstrate that distance-dependent synergy is not limited to highly context-dependent DHSs and thus constitutes a broad feature of neighboring enhancers.

We then assessed synergy between *Sox2* 19 and *Sox2* 24, another pair of DHSs that exhibit strong synergy when placed adjacent (**Fig. S8a-d**) but that are spaced 23 kb apart in the genome. We used PCR to generate a series of novel fragments from SynHPRT1R. In addition, we generated a spacer library from a human genomic BAC (RP11-158F22) encompassing a gene desert which represents a more natural but regulatory-depleted genomic region (**Fig. S7c**). The data from both spacer libraries showed a similar strong distance decay (**Fig. S9a**), confirming that dis-tance-dependent synergy is not an artifact of any single spacer library. As expected, some individual fragments deviated from the overall trend. Overall, across all spacer fragment sources, SynHPRT1R^noCpG^ showed the least deviation from regression lines, while RP11-158F22 showed the most (**Fig. S10**). Some of this deviation could be attributed to functional elements within RP11-158F22: outliers such as AvrII_6147 and AvrII_8003 overlapped DHSs in other tissues, while AvrII_642, AvrII_1755, and AvrII_3094 overlapped LINE elements (**Fig. S9a**, **Fig. S7c**). Thus, although individual spacer fragments deviated from the fitted regression lines, an overall distance-dependent synergy decay was a feature of multiple independent spacer libraries sourced from both synthetic and genomic DNA.

To enable scalable testing of multiple DHS pairs in a constant context, we developed a simplified spacer series using a discrete set of fragments. We selected four SynHPRT1R^noCpG^-derived fragments of defined lengths (231, 1159, 1874, and 4837 bp) (**Fig. S7b**) that had not previously shown orientation dependence or cryptic regulatory activity, and which spanned the distance range where interaction constraints were observed. These fragments were PCR-amplified and cloned as a spacer panel providing a reduced yet representative range of distances. This approach allowed us to efficiently assess distance-dependent DHS interaction across additional pairs, including ectopic combinations such as *Sox2* 23 with *Prdm14* +52, and *Sox2* 24 with *Sall1* +52&+53 (**Fig. S9b**). Analysis of regression fits showed that DHSs combinations respond differently to changes in inter-DHS spacing. While some pairs, such as *Sox2* 24 & 26 or *Sox2* 24 & *Sall1*+52&+53, retained high levels of activity across a broader range of distances, others, like *Sox2* 19 & 24 showed a drop in activity at shorter distances, indicating a more stringent spatial requirement for the observed synergy (**Fig. 5e**). Despite differences in enhancer identity and spacer sequence composition, all experiments recovered a similar monotonic decay in DHS synergy with increasing spacing, suggesting that while the magnitude and sensitivity to spacing varies across different DHS combinations, genomic distance broadly constrains the functional interaction of individual DHSs over the kilobase range scale.

### DHS synergy relies on a highly structured TF binding pattern

To assess specific TF site contributions to DHS synergy, we considered a series of TF recognition motif matches within *Sox2* 24 numbered 24.1–24.4 shown to be important for the function of *Sox2* 24 through deletion analysis (Brosh et al. 2023). These sites were predicted to be occupied by key mESC regulators, including the ESRR family, nuclear receptors, SOX, and forkhead proteins (**Fig. 6a**). We reengineered each deletion in the context of the *Sox2* 23 & 24 DHS pair, and delivered them in place of the *Sox2* LCR. Deletion of different TF sites showed a wide range of impact on *Sox2* activation (**Fig. 6b**). Comparing their effects on *Sox2* expression across both contexts revealed a strong correlation: deletions that most reduced expression in the *Sox2* 24-alone context, had a proportionally larger effect in the *Sox2* 23 & 24 background (**Fig. 6c**-**d**). The positive correlation indicates that while the relative contribution of each TF motif is preserved, their impact is amplified in the higher-activity *Sox2* 23 & 24 context. This suggests there is no distinction between TF sites that affect activity from those affecting synergy.

**Fig. 6.**
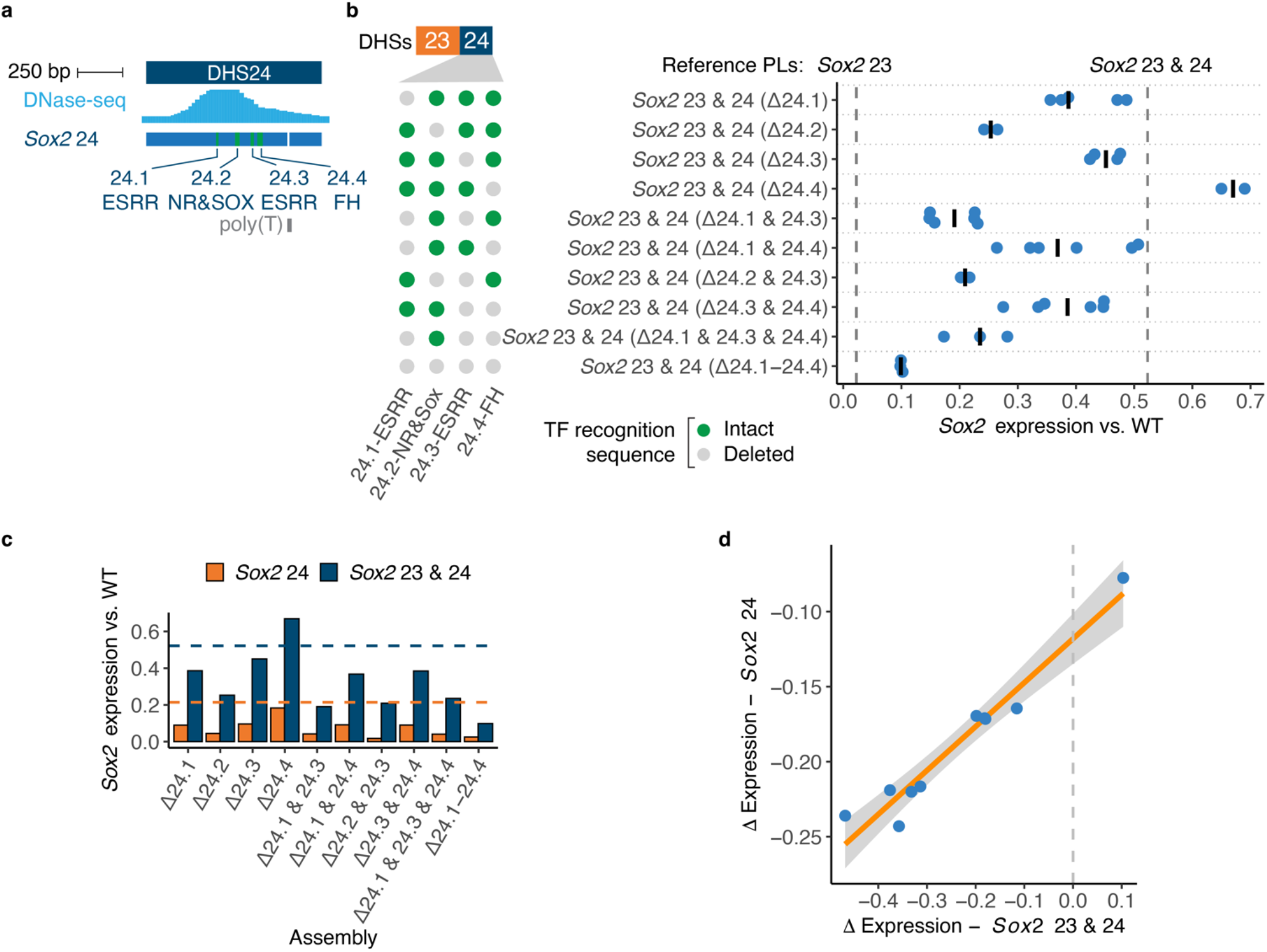
Individual TF contributions are preserved across different enhancer contexts. **a**. Detail of *Sox2* 24, showing DNase-seq data in mESC. Below, schematic representation of *Sox2* 24 payload boundaries and annotated TF motif positions. **b.** Analysis of *Sox2* 24 TF site deletions within the context of *Sox2* 23 & 24. The presence (green dots) or absence (gray dots) of each TF site is indicated. Each point represents *Sox2* expression from the engineered allele in an independent mESC clone. Bars indicate median. Expression from the BL6 allele was normalized to the CAST allele and scaled between 0 (Δ*Sox2*) to 1 (WT LCR). Vertical dashed lines indicate median expression of baseline payloads. **c**. Bar plot comparing the effect on *Sox2* expression of each TF motif deletion in the context of *Sox2* 24 (orange) (Brosh et al. 2023) and *Sox2* 23 & 24 (blue). Horizontal dashed lines indicate median expression of the corresponding baseline payloads. **d**. Comparison of the effect of matched TF motif deletions in the context of *Sox2* 24 (Brosh et al. 2023) and *Sox2* 23 & 24, suggesting that individual TF contributions are preserved between both contexts.

Expanding on these results, we next characterized two additional context-dependent DHSs: *Sox2* 19 and *Sox2* 20 (**Fig. S8a**), previously shown to synergize with *Sox2* 25–26 (Brosh et al. 2023). To examine their architecture and function more closely, we tested the activity of *Sox2* 19 and *Sox2* 20 with *Sox2* 23 and *Sox2* 24. The combination of *Sox2* 19 and *Sox2* 20 with *Sox2* 24, drove 66% of wild-type *Sox2* expression (**Fig. S8b-d**). Three context-dependent DHSs which each had little activity alone (*Sox2* 19, *Sox2* 20, and *Sox2* 23) drove significant expression when combined. Similarly, linking two copies of *Sox2* 23 yielded higher activity than a single copy (**Fig. S2**). These results together suggest that a sufficient quantity of context-dependent DHSs can approach the activity of autonomous DHSs.

We then attempted to systematically assesses the contribution of predicted TF binding sites to synergy among context-dependent DHSs. *Sox2* 19 covers only 503 bp and includes three TF sites identified through TF motif analysis and ChIP-nexus data (**Fig. S8a**). Delivery of a synthetic payload containing targeted deletions of these sites confirmed their activity and showed that deletion of the three identified TF sites within *Sox2* 19 was sufficient to completely ablate synergy with *Sox2* 24 (**Fig. S8e**). We next focused on *Sox2* 23, a longer and more complex element than *Sox2* 19. Based on TF motif analysis and ChIP-nexus data, we had previously identified TF recognition sites in *Sox2* 23, numbered 23.1–23.8 (Brosh et al. 2023). These sites were predicted to be occupied by key mESC regulators, including ZIC3, the ESRR family, forkhead proteins, and the TBX family (**Fig. 7a**). Analysis of deletion payloads showed that *Sox2* 23 is a composite element with multiple regions contributing to the interaction with *Sox2* 24 (**Fig. 7b**). Notably, even after deletion of all identified motifs in *Sox2* 23 (Δ23.1–23.8), expression of the DHS pair remained higher than with *Sox2* 24 alone, suggesting that *Sox2* 23 retains residual regulatory activity beyond the TF recognition sites mapped here. Thus, while some DHS contributions can be mapped to discrete TF sites, further work is needed to fully ascribe DHS synergy to individual TF sites.

**Fig. 7.**
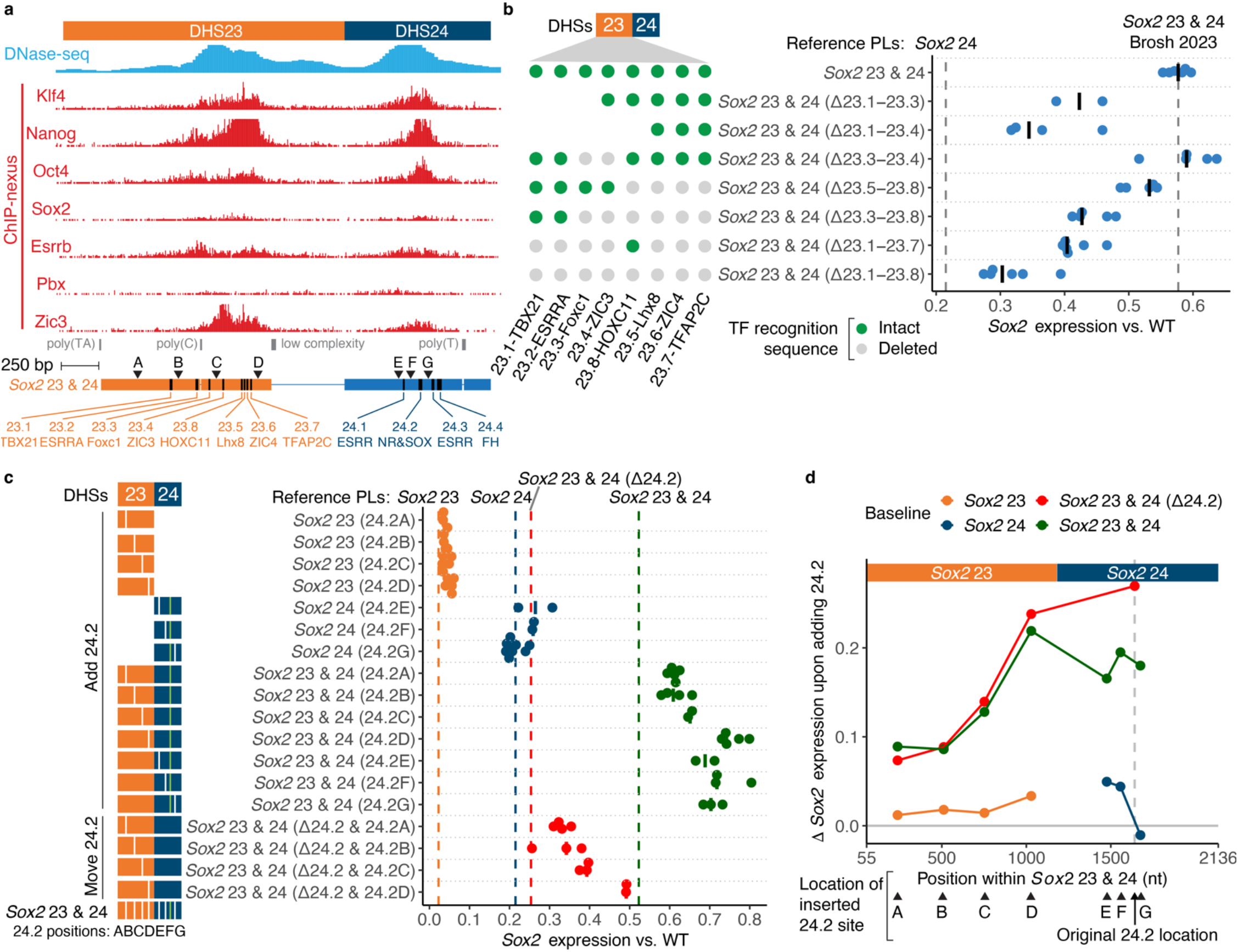
Synergy between *Sox2* 23 and *Sox2* 24 relies on a highly structured TF binding pattern. **a**. Detail of *Sox2* 23 and *Sox2* 24, showing DNase-seq and ChIP-nexus data for TF occupancy in mESC. Below, schematic representation of *Sox2* 23 & 24 payload boundaries, annotated TF motif positions, and the relative positions A-G used for TF motif 24.2 relocation. **b.** Dissection of *Sox2* 23 TF recognition sites. The presence (green dots) or absence (gray dots) of each TF site is indicated for each payload; truncations are indicated by the absence of dots. Each point represents *Sox2* expression from the engineered allele in an independent mESC clone. Bars indicate median. Expression from the BL6 allele was normalized to the CAST allele and scaled between 0 (Δ*Sox2*) to 1 (WT LCR). Vertical dashed lines indicate median expression of baseline payloads. *Sox2* 23 & 24, *Sox2* 23 & 24 (Δ23.1-23.7) and *Sox2* 23 & 24 (Δ23.1-23.8) were previously reported (Brosh et al. 2023). **c-d**. Contribution of TF site 24.2 by position within different contexts. Relocation (red) or addition (orange, blue, or green) of TF site 24.2 to *Sox2* 23, *Sox2* 24, or *Sox2* 23 & 24 payloads at positions A-G (triangles in **a**). **c**. *Sox2* expression analysis. The DHS schematic (left) illustrates the presence or absence of the original 24.2 motif (green ticks) and the position of the newly inserted copy (white ticks). **d**. Δ*Sox2* expression was computed as the mean *Sox2* expression difference between payload pairs differing solely by the presence of TF site 24.2. Vertical gray line indicates original TF site 24.2 location.

We next asked whether the overall synergy between *Sox2* 23 and *Sox2* 24 depends on a specific arrangement of TF recognition sequences across both elements. Previous deletion analysis had identified the TF site 24.2, which overlaps nuclear receptor and SOX motif matches, as the single most important TF site in *Sox2* 24 (Brosh et al. 2023). We therefore inserted a copy of TF site 24.2 at 6 different positions along the sequence of *Sox2* 23 or *Sox2* 24 (labelled A-G in **Fig. 7a**). Adding TF site 24.2 to *Sox2* 23 alone was insufficient to convert it to an autonomous enhancer like *Sox2* 24 (**Fig. 7c**, orange points). Similarly, adding a second copy of TF site 24.2 to *Sox2* 24 (blue points) did not increase activity, demonstrating that a single copy of TF site 24.2 does not limit the activity of *Sox2* 24 when on its own. In contrast, when *Sox2* 23 and 24 were delivered together, both insertion of a second copy of TF site 24.2 (green points) and its relocation (red points) increased activity above baseline. Furthermore, its contribution was strongest within 600 bp of the native TF site 24.2, and declined steeply beyond that (**Fig. 7d**). These results suggest that context-dependent enhancers are not easily made autonomous, and that context is shaped by their precise spatial configuration, highlighting a highly structured regulatory grammar underlying DHS synergy.

### Trait-associated variants reside within complex regulatory environments

Human disease- and trait-associated variation resides in regulatory DNA, but individual variants are typically analyzed independently of each other. Thus, we asked how frequently disease- and trait-associated variants reside in regulatory environments susceptible to enhancer context dependence. We investigated a uniformly processed series of genome-wide association studies (GWAS) from UK Biobank and international consortia. These included GWAS for quantitative traits including systolic and diastolic blood pressure, calcium, direct bilirubin, glucose, low-density lipoprotein (LDL) cholesterol, and triglyceride levels, red blood cell count (RBC), and diseases including hypothyroidism and type 2 diabetes (Forgetta et al. 2022). GWAS results for these traits and diseases were previously fine mapped with FINEMAP while permitting a high number of causal single nucleotide polymorphisms (SNPs) (k=20) in each configuration (Benner et al. 2016).

Since our results showed that DHS synergy extends predominantly within a 4-kb range (**Fig. 5**), we examined the DNase signal in trait-matched samples within ±4 kb windows around each trait-associated SNPs (**Fig. 8a**, **Table S8**). This analysis revealed that trait-associated variants are frequently embedded within complex regions which include multiple regulatory elements with high chromatin accessibility (**Fig. 8b**). Between 57–95% of trait-associated SNP overlapping a DHS are commonly in close proximity (<4 kb) to at least one other DHS (**Fig. 8c**). Together, these results suggest that a high proportion of trait-associated variants reside within complex clustered regulatory environments, where they may influence not only the activity of the local DHS but also interactions or synergistic effects with nearby regulatory elements. These findings underscore the importance of interpreting noncoding variants within their broader regulatory context rather than as isolated features.

**Fig. 8.**
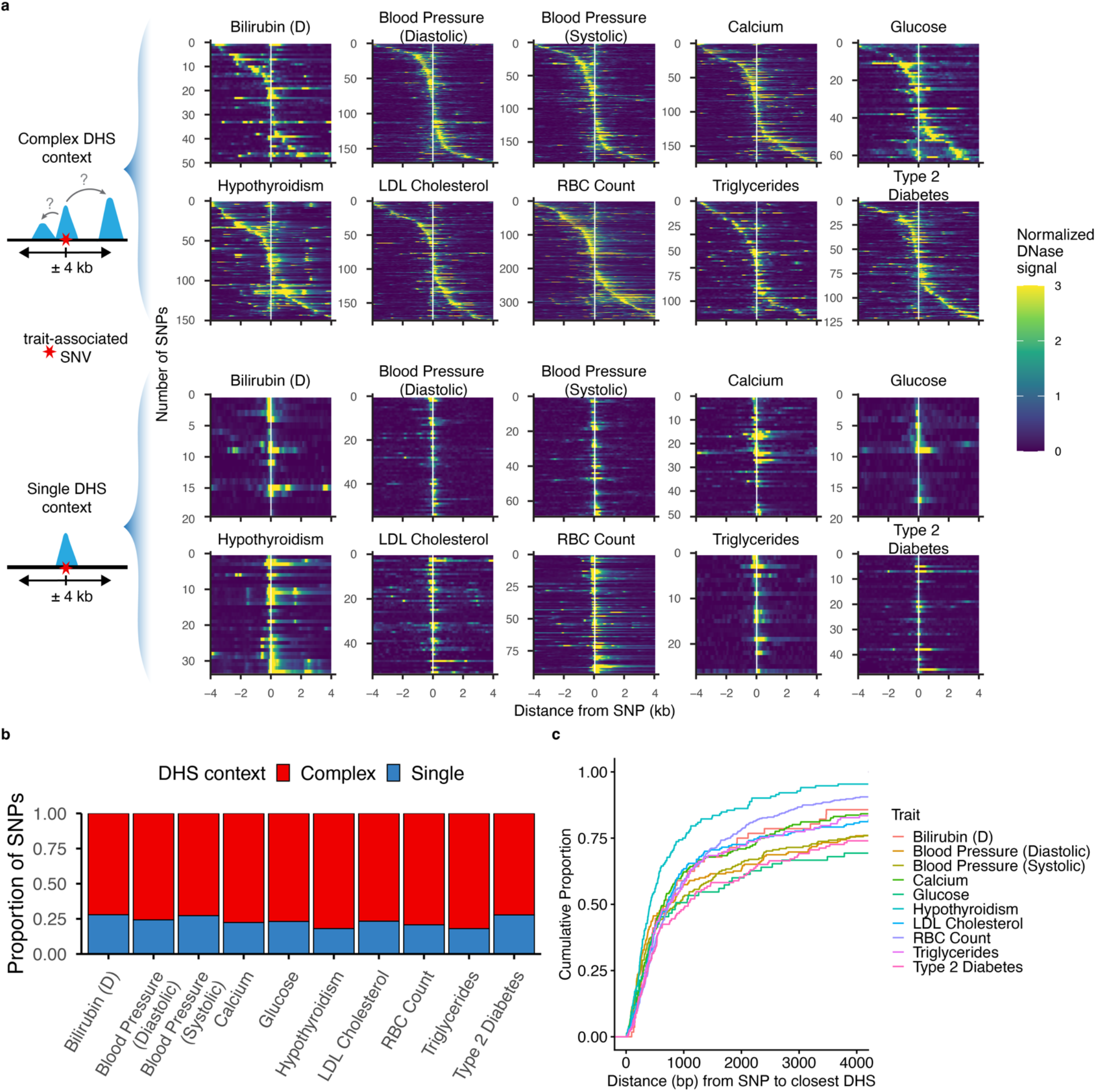
Trait-associated polymorphisms reside within complex regulatory environments. **a.** Heatmaps showing DNase-seq signal in ±4 kb windows centered on trait-associated single nucleotide polymorphisms (SNPs) overlapping a DHS (identified as DNase hotspots at 5% FDR). Each row represents an individual SNP, and heatmaps are grouped by trait across 10 phenotypes. SNP-trait-sample associations were obtained from (Forgetta et al. 2022). Bilirubin (D), direct bilirubin; RBC, red blood cell; LDL, low-density lipoprotein. DNase read density was normalized to 1M reads. **b.** Proportion of SNPs classified as residing in either complex DHS or single DHS regulatory contexts, based on the presence of additional accessibility signal within a ±4 kb window (see **Methods** for classification criteria). **c.** Cumulative proportion of SNPs as a function of distance to the nearest non-overlapping DHS.

## DISCUSSION

Enhancer clusters are key units of transcriptional regulation, but the principles governing the interactions among their constituent elements remain poorly understood. We have developed a large-scale experimental approach for functional dissection of enhancer synergy. By testing multiple heterotypic DHS pairs, we found that roughly half of them displayed some form of interaction (positive or negative synergy). This synergy was highly dependent on the identity, arrangement, and distance between the DHSs. Our data showing DHS synergy at both the *Sox2* and *Igf2/H19* loci and three different target genes indicates that DHS synergy is likely to be a widespread phenomenon rather than restricted to a single regulatory architecture. Although we inspected several hallmarks of regulatory activity for each candidate DHSs (including TF occupancy, histone modifications, or nascent transcripts), none of these alone predicted enhancer activity or synergy in our system. Some DHSs with robust reporter activity (such as *Sall1* +39&+40 or *Sall1* +27&+28) failed to drive *Sox2* expression when placed alone at *Sox2*, showing that the classical definition of an enhancer does not fully encompass their behavior at complex loci. We expect that patterns will emerge with screening of additional candidates, loci, and cell types, investigation of additional genomic parameters beyond enhancer spacing, and profiling of more detailed molecular readouts such as long-read single-molecule footprinting (Stergachis et al. 2020; Swanson et al. 2025).

A key question is whether DHS synergy is driven qualitatively by specialized TFs, or if instead synergy emerges from the quantitative accumulation of diverse transcriptional activators. While *Sox2* 19, *Sox2* 20, and *Sox2* 23 alone have little activity individually, combinations of multiple context-dependent DHSs collectively approached the activity of autonomous enhancers, fitting with past work that identified a range of context dependence rather than a strict dichotomy (Brosh et al. 2023). Both context-dependent and autonomous DHSs showed distance-dependent synergy. The effect size of individual TF motif deletions within *Sox2* 24 were strongly correlated across the *Sox2* 24 and *Sox2* 23 & 24 contexts. Finally, attempts to convert *Sox2* 23 into an autonomous enhancer by inserting the key TF site 24.2 at multiple positions failed to confer autonomous activity. Thus, our work suggests that context-dependent DHSs differ from autonomous DHSs quantitatively rather than qualitatively.

Regulatory activity fundamentally relies on proximity, making genomic distance a key parameter in cis-regulatory function. The influence of distance spans multiple distinct scales: from promoter-enhancer communication and locus architecture across tens to hundreds of kilobases, to the organization of enhancer clusters at the kilobase scale, and down to the precise arrangement of TF recognition sequences at base pair resolution within individual regulatory elements. While the impact of enhancer-promoter distance is well-recognized (Zuin et al. 2022; Thomas et al. 2025; Jensen et al. 2025), there has been less attention on the spacing between enhancers on the order of hundreds to thousands of base pairs (Small et al. 1993). It remains an open question what mechanism might underlie synergy on the order of 4 kb, whether mediated by nucleosomes (Mirny 2010) or diffusion (Karr et al. 2022). Single-molecule methyltransferase footprinting has identified distance-dependent correlation in accessibility among DHSs on a similar kilobase scale (Stergachis et al. 2020), as has analysis of allelic imbalance in bulk DNase-seq data (Maurano et al. 2015; Halow et al. 2021). Our results highlight the importance of incorporating distance effects into cis-regulatory models. Our work further supports the notion that enhancer cluster spatial organization reflects a functional arrangement that contributes to the precision, robustness, and cell-type specificity of gene regulation.

Beyond a conceptual framework which scaffolds genomic elements on distance, both the position and orientation of *Sox2* DHSs influence their activity. Thus, an important limitation of our screen is that candidate DHSs were only tested in one relative arrangement, with the context DHS in a single orientation positioned upstream of the candidate DHS. But *Sox2* 23 & *Sox2* 24 behave differently when reversed (**Fig. S2**) (Brosh et al. 2023), and even larger enhancer clusters exhibit polarity (Brosh et al. 2023; Ordoñez et al. 2024; Kassouf et al. 2025). Thus, future studies will be necessary to systematically dissect how position and orientation contribute to enhancer cluster function.

Moreover, distance and orientation imply a focus on two points, but multiple functional genomic elements are in play at any given genomic locus. Thus, attempts to disentangle this confusion must also consider “position”, which implies a coordinate within a multi-dimensional landscape of genomic elements. Our premise is that a reductive and systematic genetic approach to interrogate these functional interactions along key axes can resolve confounding observations and derive mechanistic understanding of genomic regulatory architecture.

The functional relationship of genetic variation on enhancer interactions remains largely unexplored. Our results suggest that trait-associated SNPs within these clustered regulatory neighborhoods may modulate not only local enhancer activity, but also synergistic interactions between neighboring elements. This is consistent with reports of multiple causal variants underpinning complex trait architecture (Corradin et al. 2014; Sobreira et al. 2021; Long et al. 2025). Future studies may help resolve how natural genetic variation perturbs cooperative enhancer activity across both disease-associated and normal human genetic variation.

## ACKNOWLEDGEMENTS

We thank Naydelin Zamora for technical assistance with DNA assembly. This work was partially funded by National Institutes of Health (NIH) grants RM1HG009491 (J.D.B. and M.T.M) and R01MH136353 (M.T.M.) and an Irma T. Hirschl/Monique Weill-Caulier Research Award (M.T.M.). R.B., J.D.B., and M.T.M. are listed as an inventor on a patent application describing Big-IN.

## AUTHOR CONTRIBUTIONS

R.O., G.E., R.B., and M.T.M. designed experiments. R.O., G.E., N.B.Z., and R.B. performed experiments. B.C. and J.D.B. contributed reagents. R.O., C.M., G.E., N.B.Z., and R.B. assembled DNA payloads. C.M., G.E., H.J.A., M. M, and K.B. performed sequencing. R.O., A.M.R.-d.-S., and M.T.M. performed computational analyses. R.O. and M.T.M. wrote the manuscript.

## METHODS

### Generation of barcoded payloads, delivery, and amplicon library preparation

A minimal reporter payload consisting of an EF1α promoter-GFP cassette flanked by loxM and loxP sites was constructed and subsequently barcoded prior to delivery into mESC landing pad cells (**Fig. S1a**). The reporter payload plasmid was PCR amplified using primers oRO_297 and oRO_298 (**Table S9**), treated with DpnI (R0176S) for 1 h at 37 °C, and gel-purified from a 1% agarose gel using the Zymo Clean Gel DNA Recovery Kit (Zymo Research D4001). Gibson Assembly was then performed with the amplified payload and a synthesized DNA fragment (GibsonBC, **Table S9**) according to the manufacturer’s instructions (NEB E2611L). The resulting barcoded plasmid library was purified using the Zymo Clean & Concentrate-5 kit (Zymo Research, D4014), digested with NcoI-HF (NEB R3193S) to eliminate unbarcoded plasmids (the restriction site is replaced by the barcode sequence), and purified again prior to transformation. The purified library was electroporated into MegaX DH10B-T1 electrocompetent cells (Thermo Fisher Scientific C640003) using an Eppendorf 2510 electroporator set to 1800 V. Following 1 h recovery at 37 °C, cultures were expanded in 50 mL LB containing 100 μg/mL ampicillin and grown for 16 h at 37 °C with shaking at 220 rpm. Plasmid library DNA was isolated using the ZymoPURE II Midiprep kit (Zymo Research D4201) and quantified by NanoDrop. Barcoded payload library delivery was performed as described below.

To assess the number of uniquely barcoded, on-target payload integrations achieved after delivery, amplicon libraries were generated from genomic DNA extracted from payload-delivered cells, as previously described (Ribeiro-Dos-Santos et al. 2022) with minor modifications. A single-cycle PCR (using one primer binding the payload and one primer binding the flanking genome) was first performed to incorporate unique molecular identifiers (UMIs) and the internal portion of the P5 adapter. Eight replicate 50 μL reactions were prepared per sample, each containing 500 ng genomic DNA, 1× KAPA HiFi HotStart ReadyMix (Kapa Biosystems, KK2602), and 200 nM primer oRO_598 (*Igf2* deliveries) or oRO_600 (*Hprt* deliveries) (see primer sequences in **Table S9**). Cycling conditions were 95 °C for 5 min, 60 °C for 1 min, and 72 °C for 10 min. Replicates were pooled and purified using 1.8x 18% Sera-Mag magnetic beads (Cytiva, 65152105050250) in polyethylene glycol, eluting in 20 μL. Libraries were then amplified in two nested PCR steps to complete Illumina adapter addition. In the first round, inner P5 and P7 adapter sequences were added using oRO_280 (*Igf2* deliveries) or oRO_279 (*Hprt* deliveries) as forward primers and P7_Halfsite_Short as reverse primer (see primer sequences in **Table S9**). Reactions contained 1 μL purified DNA, 1× KAPA HiFi HotStart ReadyMix, and 200 nM of each primer in a 50 μL final volume. Cycling conditions were 95 °C for 5 min; 32 cycles of 95 °C for 15 s, 65 °C for 15 s, and 72 °C for 30 s; followed by 72 °C for 10 min. Products were purified as above. In the second round, outer indexed P5 and P7 adapters (P5_amplicon_Index and P7_amplicon_Index; **Table S9**) were added. Reactions contained 1 μL DNA purified from the previous PCR, 1× KAPA HiFi HotStart ReadyMix, and 200 nM of each primer in a 50 μL final volume. Thermal cycling conditions were: 95 °C for 5 min; 8-10 cycles of 95 °C for 15 s, 71 °C for 15 s, and 72 °C for 30 s; followed by 72 °C for 10 min. Final libraries were purified using 1.8x 18% Sera-Mag magnetic beads (Cytiva, 65152105050250) in polyethylene glycol, eluted in 20 μL, and quantified using the Qubit dsDNA HS assay (Thermo Fisher Scientific, Q32851). Libraries were sequenced on an Illumina NextSeq 500, and barcode diversity was quantified using a previously described pipeline (Ribeiro-Dos-Santos et al. 2022). Only barcodes supported by >10 UMIs were included in downstream analyses.

### Payload assembly

Different strategies were used to assemble payloads into entry vectors (**Table S1**, **Table S2**, and **Data S1**), using a collection of oligos and synthetic DNA fragments (**Table S9**):

#### 1) Golden Gate Assembly

DNA fragments were assembled into three different entry vectors: pLM1110 (Brosh et al. 2023), pRB051 (a derivative of pLM1110) (Brosh et al. 2023), or pINEAPLE (Addgene #250295). pINEAPLE was cloned by performing BsaI Golden Gate Assembly of the synthetic fragment qRB_005 (**Table S9**) into pAV10 (Addgene #63213). Payloads were assembled with Golden Gate Assembly as previously described (Brosh et al. 2023), briefly 100 ng of entry vector was mixed with 20 ng of each DNA fragment, 0.4 μL of the appropriate restriction enzymes, either Esp3I (NEB R0734S) or BsaI-HFv2 (NEB R3733S), 1.5 μL 1 mg/mL BSA, 1 μL T4 Quick Ligase (NEB M2200S) and 1.5 μL T4 Ligase Buffer (NEB) in a total volume of 15 μL. Reactions were cycled 40 times between 37 °C (5 min) and 16 °C (4 min), heat inactivated at 60 °C (5 min) and 80 °C (5 min) before transforming into TransforMax EPI300 cells for pLM1110 or pRB051-derived payloads and JM109 for pINEAPLE-derived payloads.

#### 2) Restriction enzyme cloning

Spacer fragments were cloned into source payloads using restriction enzyme-based cloning. Briefly, source payload vectors were digested with BsiWI-HF (NEB R3553S) or AvrII (NEB R0174S) in a reaction containing 1× NEB CutSmart buffer at 37 °C for 1 h, with Quick CIP (NEB M0525S) included to prevent vector recircularization. PCR-amplified spacer fragments and BAC DNA were digested with the same restriction enzyme. BAC DNA were additionally digested with NotI-HF (NEB R3189S) or I-SceI (NEB R0694L) to prevent incorporation of backbone fragments. Digested fragments were purified using 0.8x 18% Sera-Mag Magnetic Beads (Cytiva 65152105050250) in polyethylene glycol. Purified inserts were ligated into the digested vector using T4 DNA ligase (NEB M0202S) at 16 °C overnight. Ligated products were transformed into *E. coli* JM109 by heat shock at 42 °C for 45 seconds, followed by recovery in SOC medium at 37 °C for 1 h. Bacterial cultures were expanded in 100 mL LB medium overnight at 37 °C, and plasmid DNA was extracted using the ZymoPURE II Plasmid Midiprep Kit (D4201). Cloned constructs were verified by short-read sequencing to confirm pool representation.

#### 3) Yeast assembly

First, an entry vector was constructed in *E. coli* using Golden Gate Assembly (**Table S1**). This vector carried two 500-bp homology arms corresponding to the desired payload flanks. A unique restriction site was included between the homology arms to allow for in vitro linearization. Vectors were linearized by overnight digestion with BsiWI-HF (NEB R3553S) at 37 °C, followed by gel purification using the Zymoclean Gel DNA Recovery Kit (Zymo Research D4002). Yeast cultures were then transformed with 50 ng of digested entry vector and 1 µg of source BAC DNA containing the target sequence using the lithium acetate method, and transformants were selected on SC -LEU medium. For screening, DNA was isolated from individual colonies by resuspension in 10–40 µL of 20 mM NaOH, followed by three cycles of boiling at 95 °C for 3 min and cooling at 4 °C for 1 min. PCR reactions used 10 µL of GoTaq Green Master Mix (Promega M7123) with 2 µL of lysate as template and 0.5 µM primers targeting the newly formed junctions. Candidate clones were further transferred into bacteria by extracting the payload DNA using the Zymo Yeast Miniprep I Kit (Zymo Research D2001) and electroporating it into *E. coli* TransforMax EPI300 cells (Lucigen EC300150) according to the manufacturer’s instructions.

### Payload DNA screening, preparation, and sequencing verification

For initial verification, each payload DNA was prepped differently according to the backbone replication conditions. pLM1110 and pRB051-derived payloads, including an inducible copy number backbone, were picked into 5 mL LB with kanamycin (50 μg/mL) supplemented with 1X Copy-Control Induction Solution (Lucigen CCIS125) and cultured overnight at 30 °C with shaking at 220 RPM. Payload DNAs were isolated using the ZR BAC DNA Miniprep Kit (Zymo Research D4048). pINEAPLE-derived payloads, including a high copy number backbone, were picked into 5 mL LB with ampicillin (100 μg/mL) and cultured overnight at 37 °C with shaking at 220 RPM. Payload DNAs were isolated using the Zyppy Plasmid Miniprep Kit (Zymo Research D4020). Payload assembly verification was then performed by Sanger Sequencing, short-read, or long-read sequencing methods (**Table S1**).

Sequence-verified clones were then prepped at larger scale for delivery into mESC cells:

i. Individual payloads were grown according to their backbone replication conditions. pLM1110 and pRB051-derived payloads were picked into 2.5 mL LB with kanamycin (50 μg/mL) and grown overnight at 30 °C, then subcultured 1:100 in 250 mL of LB with kanamycin (50 μg/mL) supplemented with 1X CopyControl Induction Solution (Lucigen CCIS125) and grown for an additional 8 hours at 30 °C. pINEAPLE-derived payloads were picked into 50 mL LB with ampicillin (100 μg/mL) and grown overnight at 37 °C.
ii. For pooled deliveries of payloads cloned individually, single colonies for each payload clone were picked into 5 mL of LB with ampicillin (100 μg/mL) and cultured overnight at 30 °C with shaking at 220 RPM. To generate equimolar pools of each payload DNA, cultures were then pooled based on OD600 to 5 mL total volume and expanded in 45 mL of fresh LB with ampicillin (100 μg/mL) (1:10 dilution) for 4 h at 37 °C. Representation of each payload in the pool was further confirmed by sequencing.

For transfection, DNA was purified using the ZymoPURE II Midiprep kit (Zymo Research D4201) for DNAs <20 kb or using the Nucleobond XtraBAC kit (Takara 740436) for DNAs >20 kb. DNA preps were stored at 4 °C.

### mESC culture and engineering

C57BL6/6J x CAST/EiJ (BL6xCAST) clone 4 male mESCs49 were kindly provided by David Spector (Cold Spring Harbor Laboratory, Cold Spring Harbor, NY) and cultured as previously described (Brosh et al. 2021). Cells were maintained on 0.1% gelatin-coated plates (EMD Millipore ES006-B) in 80/20 medium comprising 80% 2i medium and 20% mESC medium. 2i medium contained a 1:1 mixture of Advanced DMEM/F12 (ThermoFisher 12634010) and Neurobasal-A (ThermoFisher 10888022) supplemented with 1% N2 Supplement (ThermoFisher 17502048), 2% B27 Supplement (ThermoFisher 17504044), 1% GlutaMAX (ThermoFisher 35050061), 1% PenStrep (ThermoFisher 15140122), 0.1 mM 2-mercaptoethanol (Sigma M3148), 1,250 U/mL LIF (Sigma ESG1107), 3 mM CHIR99021 (R&D Systems 4423), and 1 mM PD0325901 (Sigma PZ0162 and StemCell Tech #72184). mESC medium contained KnockOut DMEM (ThermoFisher 10829018) supplemented with 15% FBS (BenchMark 100-106), 0.1 mM 2-mercaptoethanol (Sigma M3148), 1% GlutaMAX (ThermoFisher 35050061), 1% MEM nonessential amino acids (ThermoFisher 11140050), 1% nucleosides (EMD Millipore ES008-D), 1% Pen-Strep (ThermoFisher 15140122), and 1,250 U/mL LIF (Sigma ESG1107). Cells were grown at 37 °C in a humidified atmosphere of 5% CO2 and passaged on average twice per week. Cells were routinely tested for mycoplasma contamination.

Payload deliveries were performed as previously described (Brosh et al. 2021) with minor adjustments based on payload size. Briefly, 1.5-5×10^6^ mESCs were transfected with 1.5-5 μg of payload DNA and 1.5-5 μg of pCAG-iCre plasmid (Addgene #89573), and plated at densities ranging from 1.5×10^5^ to 5×10^6^ cells per 10 cm plate. Following transfection, mESCs were selected with 10 mg/mL blasticidin for 2 days starting day 1 post-transfection and with 2 nM proaerolysin for 2 days starting day 7 post-transfection. Individual mESC clones were manually picked approximately 10 days post-transfection into gelatinized 96-well plates prefilled with 30 μL TrypLE-Select, which was neutralized with 200 μL mESC medium. Immediately after, 25 μL were transferred into a new plate with 75 μL of 2i media, obtaining two replica plates with 90% and 10% relative densities. Three days later, crude gDNA was extracted from the 90% plate using the Wizard SV 96 Genomic DNA Purification System (Promega A2370) for qPCR genotyping and DNA library preparation. Candidate clones were then expanded from the 10% density plate for further phenotypic characterization.

### mESC PCR genotyping

Genotyping of mESC clones was performed by real-time quantitative PCR (qPCR) as previously described (Brosh et al. 2023). Briefly, qPCR reactions were carried out in 384-well format using KAPA SYBR FAST (Kapa Biosystems KK4610) on a LightCycler 480 Real-Time PCR System (Roche). Each well was prefilled with 5 μL of KAPA SYBR FAST and 4 μL of water. Primers (50 nL of 100 μM stock) and genomic DNA (500 nL per reaction) were dispensed by the Echo 550 liquid handler system to ensure consistency. Thermal cycling conditions were: 95 °C for 3 min, then 40 cycles of 95 °C for 3 sec, 63 °C for 20 sec, and 72 °C for 20 sec. Assays targeted the expected left and right homology arm junctions, loss of the landing pad, and absence of the payload vector backbone. For payloads flanked by unique nucleotide sequences (UNS), UNS-specific primers were included. Primer sequences are listed in **Table S9**.

### High-throughput sequencing verification and deconvolution of payload identity

Illumina sequencing libraries were prepared from purified DNA using the One-Pot dsDNA protocol or the NEBNext Ultra FS II kit (NEBE7805L) as previously described (Brosh et al. 2023) (**Table S7**). Hybridization capture (Capture-seq) for targeted resequencing of engineered regions was performed as previously described (Brosh et al. 2021). Libraries were sequenced on an Illumina NextSeq 500 or 2000. Bait sets and sequencing statistics are listed in **Table S7**. Sequencing parameters and the analysis pipeline are detailed in (Brosh et al. 2023). The full processing pipeline is available at https://github.com/mauranolab/mapping.

Payloads delivered individually were verified by coverage depth analysis (**Table S10**). Coverage depth was quantified over payload vector backbone, landing pad, pCAG-iCre custom references, payload sequence, sequence within the LP region not represented in the payload, and sequence flanking the LP and targeted by Capture-seq. Coverage depth thresholds were set to require little or no coverage over custom references (LP, vector backbone, and pCAGiCre). Coverage at payload and non-payload regions was normalized against flanking regions covered by the same bait with coverage thresholds were set to 0.5±0.4 over non-payload regions, and 1±0.4 over payload regions at the delivery locus (*Sox2* or *Igf2)*. Clones with ambiguous coverage for sequences not expected to be present were evaluated by Bamintersect to exclude the presence of genomically integrated copies. Individually spacer delivered payloads were further validated by k-mer sequence analysis (see below).

Bamintersect (Brosh et al. 2021) was used to verify on-target payload integration for all deliveries, through identification of read pairs mapping across the junction between integration site and payload. Read pairs were hierarchically classified according to their mapping positions, based on a prior approach (Brosh et al. 2021). Briefly, left (LeftJx) and right (RightJx) junctions were defined as read pairs mapping within 1.5 kb of the payload start and end coordinates, respectively; endogenous read pairs were attributed to endogenous genomic copies of sequence engineered into payloads. Junctions with few reads (<1 ReadsPer10M or representing less than 1% of the total ReadsPer10M) or covering less than 75 bp were discarded. Clones were classified as PASS if a single payload-LP junction (left or right) was detected (**Table S11**).

We used several approaches to deconvolute payload identity for multiplexed delivery of payload pools. Pooled spacer payload deliveries were deconvoluted using the count_table.sh tool from bamintersect (Brosh et al. 2021) to identify junctions between flanking DHSs and internal spacer (**Table S12**). Spacer identity was identified by the presence of correctly oriented left and right junctions. For spacer payloads comprising multiple fragments resulting from partial AvrII digestion of the source BAC, spacer identity was inferred from the pattern of bamintersect-detected junctions between the constituent AvrII fragments and confirmed manually by examining coverage. Clones with no junction detected were suspected to carry no spacer or small spacers (AvrII_53) and were confirmed by k-mer sequence analysis.

For other pooled payload deliveries and spacer clones with no detected junction, we applied a k-mer analysis which quantified the presence of short unique or near-unique sequences within sequencing reads (**Table S13** and **Table S14**). K-mers were manually chosen to inform on DHS position, junctions between DHSs or spacers, and internal TF insertions or deletions. informative k-mers were assigned to each payload (**Table S15**). Clones perfectly matching the expected k-mers were assigned to the that payload, otherwise they were assigned to the nearest payload by Manhattan distance. If there were two or more nearest payloads, the clone was labeled unknown. Only k-mer hits accounting for at least 5% of the total number of unique sequence reads were considered when defining a clone profile. Only the payloads present in the delivery pool were considered.

### RNA Isolation, cDNA synthesis and mRNA expression analysis by qRT-PCR

RNA extraction, cDNA synthesis and qRT-PCR were performed as previously described (Brosh et al. 2023). Briefly, RNA was isolated using the RNeasy-mini Kit (QIAGEN 74106) and treated with Turbo DNA-free kit (ThermoFisher AM1907) following the “rigorous DNase treatment”. cDNA synthesis was performed from 1-2 µg total RNA using the High-Capacity cDNA Reverse Transcription Kit (ThermoFisher 4368814), including -RT controls for a subset of samples. Real-time quantitative reverse transcription PCR (qRT-PCR) was performed using KAPA SYBR FAST (Kapa Biosystems KK4610) on a LightCycler 480 (Roche). An Echo 550 (Beckman Coulter) liquid handler was used for loading 348-well plates as described above. Allele-specific primers (**Table S9**) were used with thermal cycling conditions as previously described for *Sox2* (Brosh et al. 2021) (95 °C for 3 min, then 40 cycles of 95 °C for 3 sec, 57 °C for 20 sec, and 72 °C for 20 sec) and *Igf2*/*H19* (Ordoñez et al. 2024) (95 °C for 3 min, then 40 cycles of 95 °C for 3 sec, 60 °C for 10 sec, and 72 °C for 10 sec). Assays were performed in triplicate wells, and replicate wells on the same plate were averaged after masking technical outliers. Raw Ct values are listed in **Data S2**. To quantify expression of *Sox2*, Ct values were normalized to *Gapdh*, and allele-specific expression was quantified using the 2^ΔCt[CAST–BL6]^ method. Fold change was scaled from 0 (Δ*Sox2*) to 1 (WT) by subtracting the mean fold change for ΔSox2 from all data points then dividing by the mean fold change of the WT LCR samples (**Table S5**). To quantify expression of *H19* and *Igf2*, Ct values were normalized to *Gapdh* and expression was reported as 2^-ΔCt^ (**Table S6**).

For spacer analyses, fragments were removed if they manifested a difference in activity between the two spacer orientations greater than 0.13 in terms of normalized *Sox2* expression.

### Candidate DHS TF motif analysis

TF motif matches overlapping each DHS were extracted using ‘`bedtools intersect`’. Motif annotations were obtained using FIMO with a cutoff of *P* < 10^-5^ (Grant et al. 2011) on published motifs derived from SELEX (Jolma et al. 2013). To reduce redundancy across similar motifs, individual motif matches were clustered by TF family using a previously published mapping (Maurano et al. 2015).

### RNA-seq data

RNA-seq transcriptional profiles from 6 mESC samples were obtained from the GEO repository (GSM6912138–GSM6912143 in GSE222038) (Camellato et al. 2024). Analysis was performed as previously described (Ordoñez et al. 2024). Briefly, reads were aligned to the mouse genome build mm10 using STAR and mapped to transcripts using RSEM (Li and Dewey 2011). Raw reads counts were normalized to FPKM using DESeq2 (Love et al. 2014).

### DNase-seq signal profiling around trait-associated SNPs

Trait-associated fine-mapped single nucleotide polymorphisms (SNPs), DNase-seq, and trait-DNase associations were obtained from (Forgetta et al. 2022). We analyzed 32,662 SNPs with FINEMAP log_10_(Bayes Factor) > 2). Fine-mapped SNPs were converted from hg19 to hg38 coordinates using UCSC liftOver. We used 31 DNase-seq datasets processed using a uniform mapping and peak calling pipeline (https://github.com/mauranolab/mapping/tree/master/dnase) with DHSs identified using hotspot2 (https://github.com/Altius/hotspot2) at a cutoff of 5% false discovery rate (FDR). We used the published associations to assign DNase-seq samples to traits (**Table S8**).

For each trait, SNPs overlapping DHSs were identified in the corresponding trait-matched DNase-seq samples. To avoid redundancy at SNPs overlapping a peak in >1 DNase sample, the sample with the highest DNase signal was selected. To analyze chromatin accessibility profiles around each SNP, DNase signal was extracted in ±4 kb windows centered on each SNP using *computeMatrix* from deepTools (Ramírez et al. 2016). Within these windows, we identified peaks as regions ≥ 150 bp where signal intensity exceeded ≥ 0.5× of the signal over the SNP. SNPs were classified as complex DHS context if either: (i) the SNP overlapped a peak extending ≥ 500 bp; or (ii) the window ±4 kb contained multiple peaks. Additionally, we computed the distance from each SNP within a DHS to the nearest non-overlapping DHS using the BEDOPS tool ‘`closest-features --dist --no-overlaps`’ (Neph et al. 2012).

## SUPPLEMENTARY MATERIALS

### SUPPLEMENTARY FIGURES

**Fig. S1.**
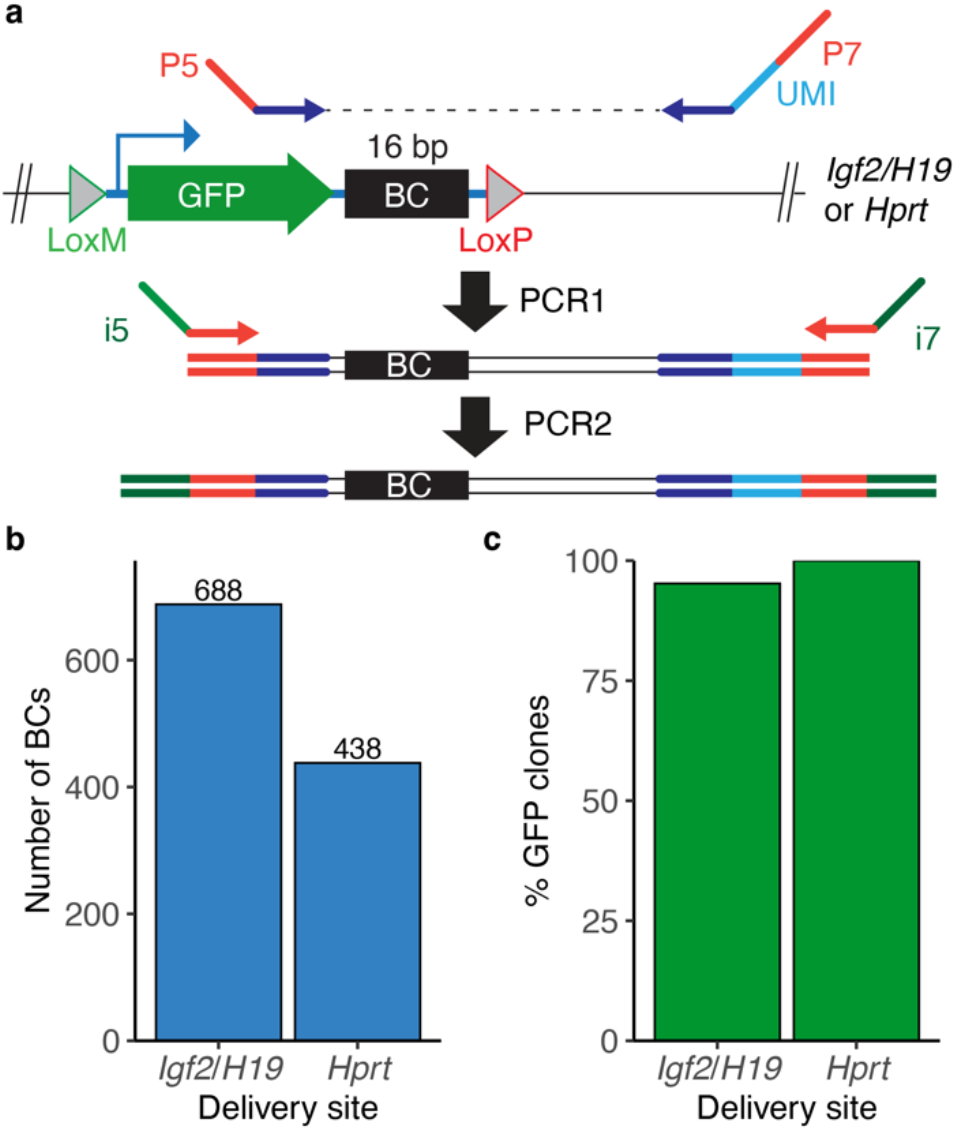
Barcode-based assessment of multiplexed Big-IN delivery efficiency. **a**. Experiment schematic including payload encoding GFP with a 16-bp barcode (BC) in its 5’UTR. The payload was delivered to two previously established mESC lines harboring Big-IN landing pads at *Igf2*/*H19* and *Hprt* (Ordoñez et al. 2024; Camellato et al. 2024). Two-stage nested PCR with an 8-bp unique molecular identifier (UMI) for single-molecule counting was used to survey barcode diversity. **b**. Number of unique barcodes identified per delivery. **c**. Percentage of surviving clones that were GFP^+^.

**Fig. S2.**
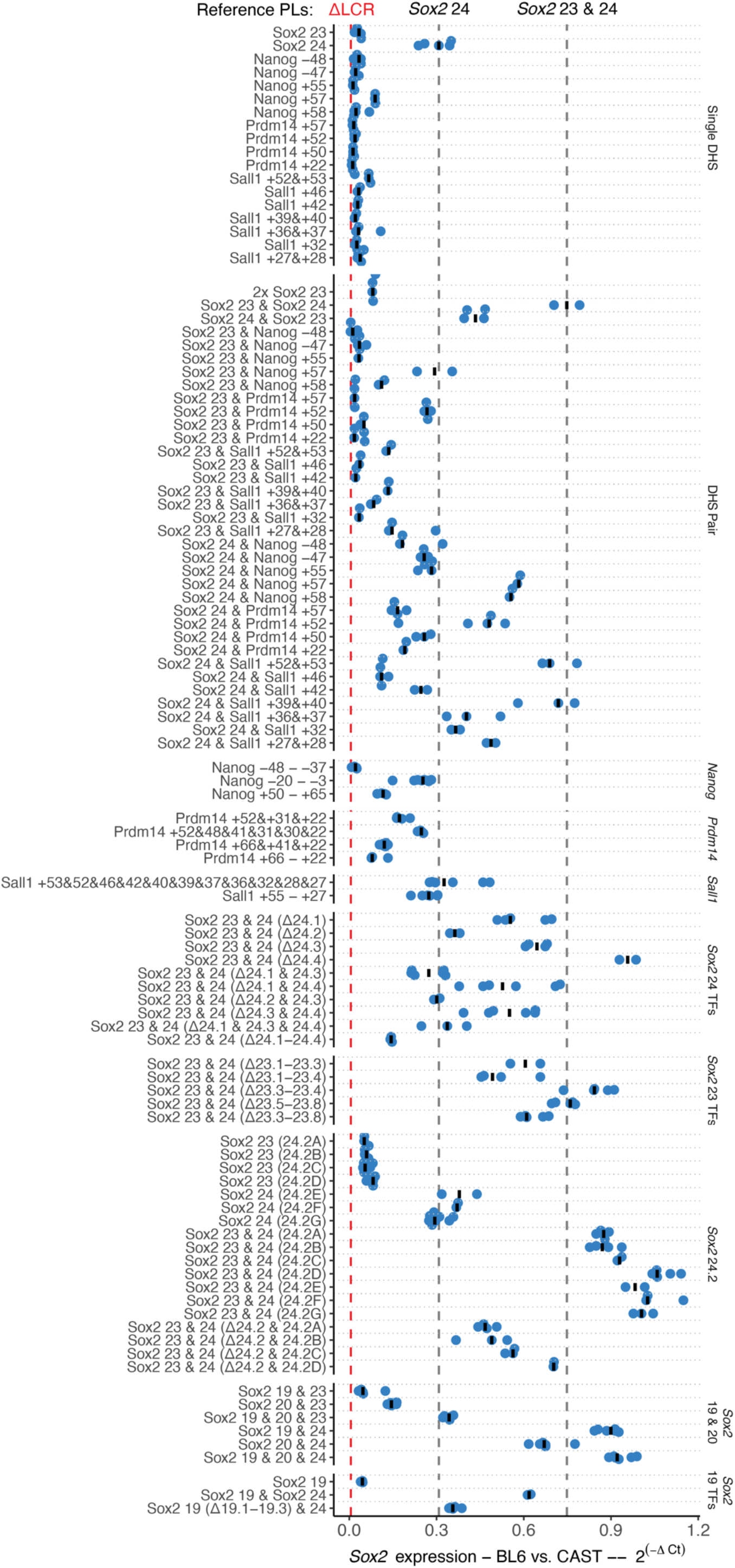
*Sox2* qRT-PCR expression values. *Sox2* expression for payloads described in this paper. Displayed is the expression of the BL6 allele of Sox2 relative to the CAST allele, calculated as 2^ΔCt[CAST-BL6]^. Points represent individual mESC clones and bars indicate median. Vertical dashed lines indicate median expression of baseline payloads. Spacer library pools are not included.

**Fig. S3.**
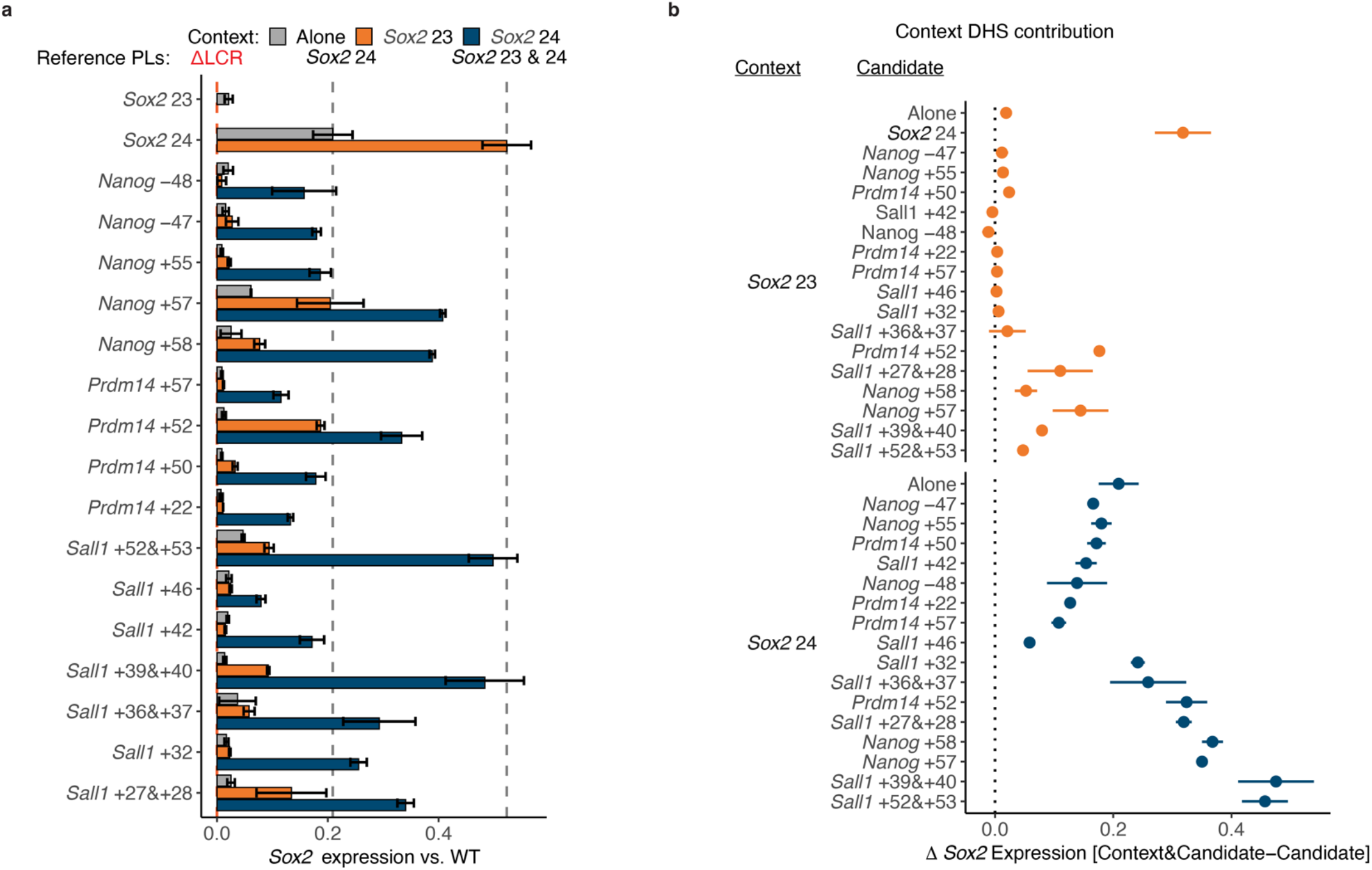
Contribution of candidates for each context DHS. **a**. Expression analysis of candidate DHSs tested alone (gray), or in combination with *Sox2* DHSs (orange or blue). Each bar represents the mean ± SD. Expression from the BL6 allele was normalized to the CAST allele and scaled between 0 (Δ*Sox2*) to 1 (WT LCR). Vertical dashed lines indicate median expression of baseline payloads. **b**. Contribution of each DHS, computed as the difference in *Sox2* expression between payload pairs differing solely by the presence of the context DHS. Points indicate mean expression change. Error bars indicate SD across all pairwise combinations of clones. Expression is scaled between 0 (Δ*Sox2*) and 1 (WT LCR).

**Fig. S4.**
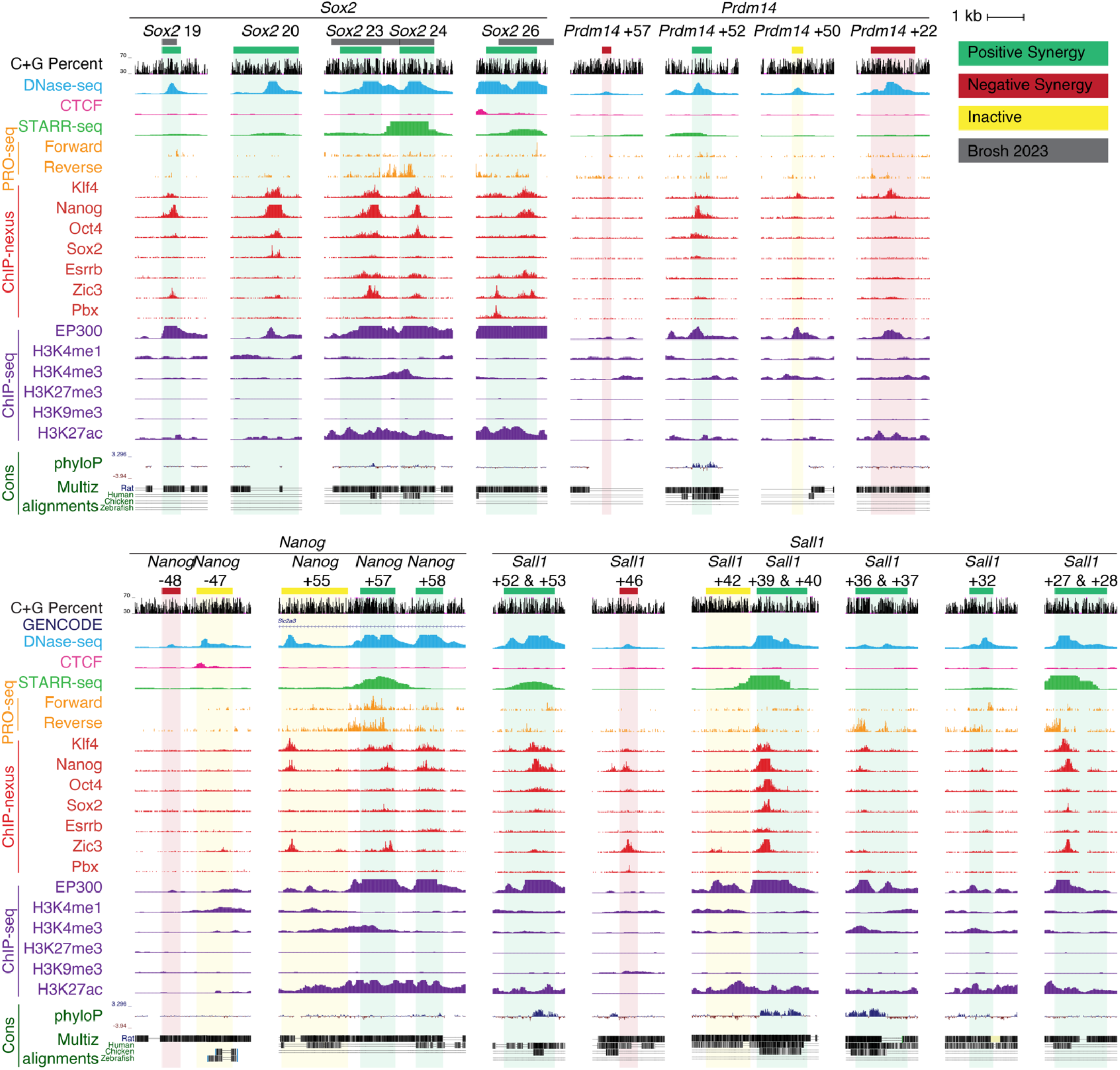
Regulatory landscape at candidate DHSs. Candidate DHSs at top are shaded in a different color according to its mode of interaction with neighboring DHSs. *Sox2* 23, *Sox2* 24, and *Sox2* 26 in this paper correspond to syn23, syn24, and syn26 previously described in (Brosh et al. 2023); other *Sox2* DHSs from that paper are shown shaded in grey; the exact DHS and coordinates used for each payload are detailed in **Table S1** and **Table S2**. Shown below are DNase-seq, CTCF ChIP-seq, STARR-seq reporter activity, PRO-seq for nascent RNA transcription, ChIP-nexus data for selected mESC regulators, EP300 and histone ChIP-seq in mESC. Sequence conservation is shown by phyloP (Placental Mammals), and Multiz Alignments with selected species. DNase-seq data for ES_CJ7 (ENCLB163SYJ, DS13320) (Vierstra et al. 2014) and ChIP-seq data (ENCLB811KZV, ENCLB074BRJ, ENCLB326DUQ, ENCLB009NHA, ENCLB738HLN, ENCLB253ATP, ENCLB187WWY) (Yue et al. 2014) were obtained from https://www.en-codeproject.org. ChIP-nexus (Avsec et al. 2021) for Zic3 (GSM4087824), Pbx (GSM4087823), Esrrb (GSM4087822), Sox2 (GSM4072777), Oct4 (GSM4072776), Nanog (GSM4072778), and Klf4 (GSM4072779), PRO-seq (Etchegaray et al. 2019) (GSE130691), and STARR-seq (Peng et al. 2020) (GSM4261634) were obtained from the GEO repository.

**Fig. S5.**
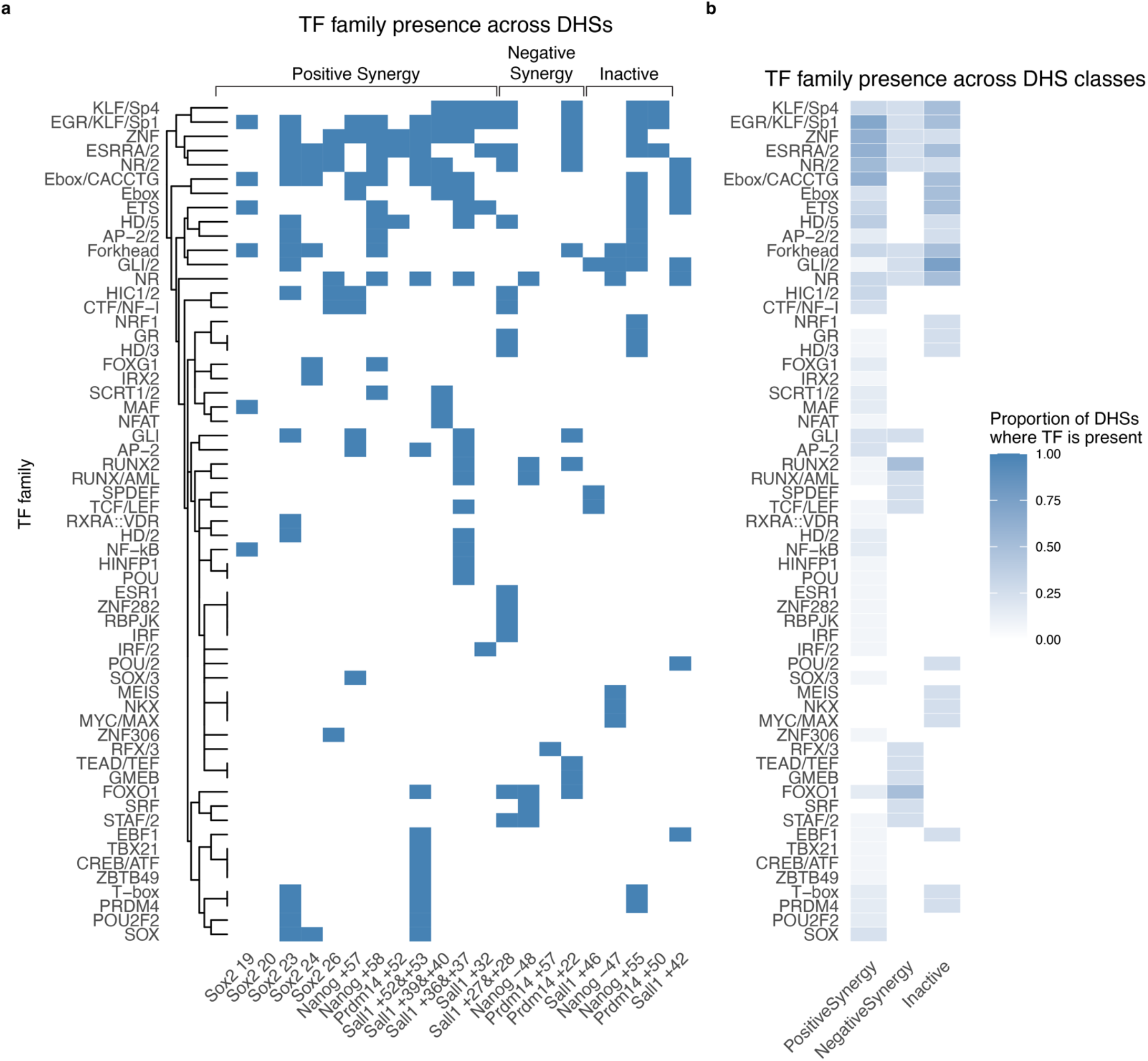
TF motif family representation at candidate DHSs.a. Presence/absence matrix of TF motif families across candidate DHSs. Each column represents a DHS, grouped by class (positive synergy, negative synergy, inactive) according to Fig. 2a, and rows indicate TF families (collapsed from individual motifs). Blue denotes presence of at least one motif from the indicated TF family within the DHS. Dendrogram represents hierarchical clustering of TF families based on Euclidean distance. **b.** Heatmap showing the proportion of DHSs within each class where a given TF family is present.

**Fig. S6.**
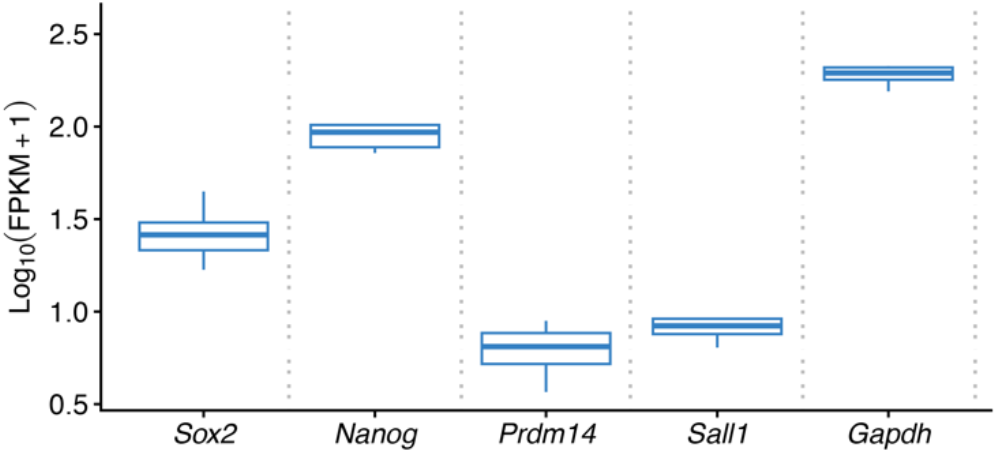
**Expression levels of enhancer cluster target genes.**Expression shown as Log_10_(FPKM+1) using published RNA-seq data (Camellato et al. 2024). Bar indicates median, hinges indicate quartiles, and whiskers indicate 1.5 times the interquartile range.

**Fig. S7.**
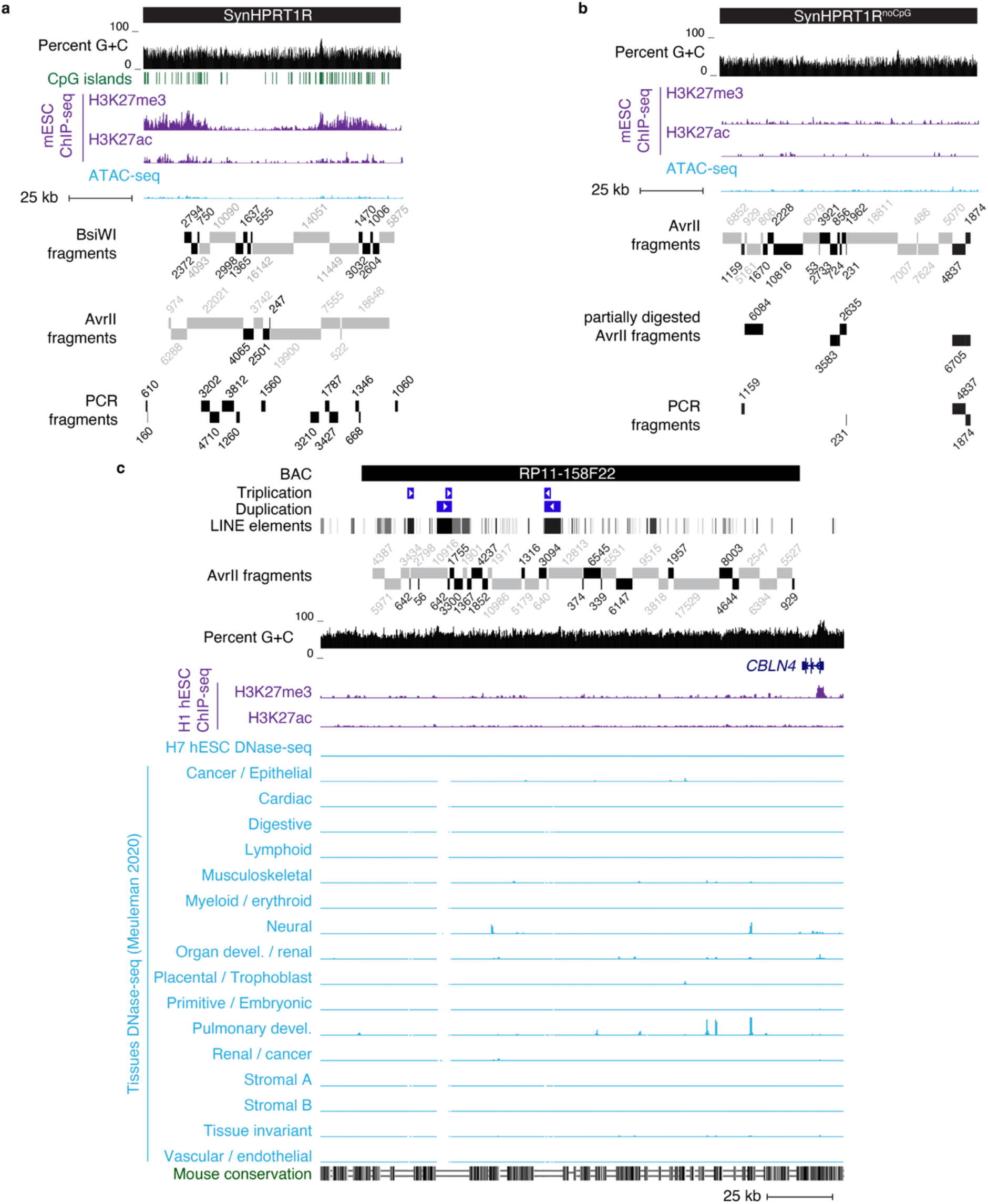
**Sequence features and chromatin profiles of spacer DNA.**Genome browser view of spacer sources. Spacer fragments detected in engineered mESC clones are highlighted in black. **a-b**. SynHPRT1R (**a**) and SynHPRT1R^noCpG^ (**b**) including H3K27me3 and H3K27ac ChIP-seq and ATAC-seq profiles from the corresponding DNA integrated at the *Sox2* locus in mESCs (Camellato et al. 2024). Shown in **a** are predicted CpG islands (Camellato et al. 2024). AvrII fragments from partial digestion are included in **b** (**Methods**). **c.** Human BAC RP11-158F22 showing RepeatMasker LINE elements and internal triplication and duplication. Shown below are H3K27me3 and H3K27ac ChIP-seq from H1 embryonic stem cells (H1 hESC) (ENCODE ENCLB788MUI and ENCLB933RET) (ENCODE Project Consortium 2012), DNase-seq data are from H7 hESC (ENCLB449ZZZ, DS11909) (Thurman et al. 2012), and meta-DNase-seq tracks from non-negative matrix factorization (Meuleman et al. 2020). Mouse conservation represents MultiZ alignment.

**Fig. S8.**
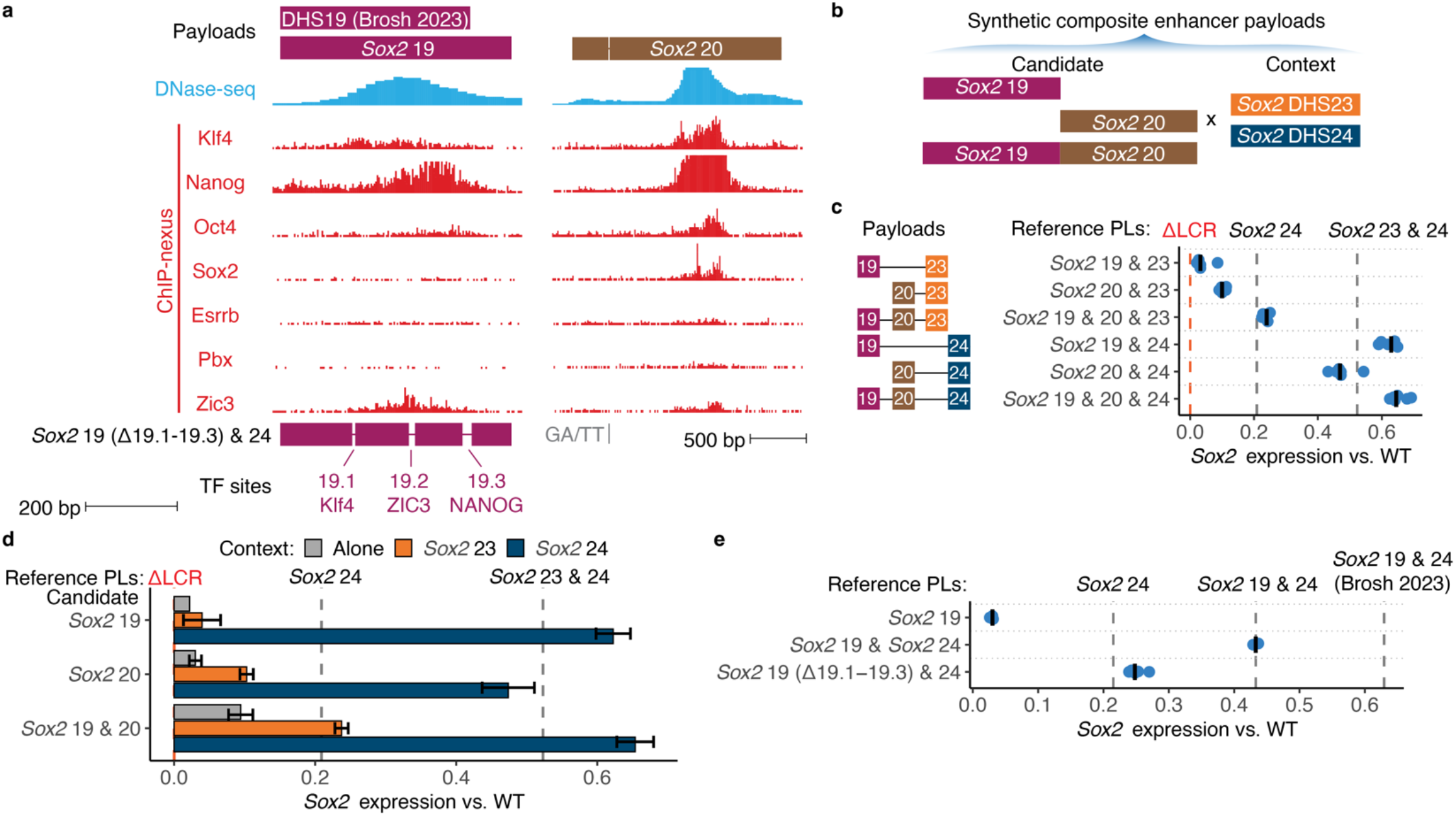
Activity and context sensitivity of *Sox2* 19 and *Sox2* 20. **a.** Detail of *Sox2* 19 and *Sox2* 20. Shown below are DNase-seq and ChIP-nexus data in mESC. Schematic below represents *Sox2* 19 Δ19.1-19.3 payload including TF site deletions. GA/TT indicates a mutation in *Sox2* 20 to remove an internal Esp3I site. *Sox2* 19 payloads in **b-d** use boundaries from (Brosh et al. 2023), payloads in **e** use extended boundaries to capture larger region. **b.** Schematic of strategy to identify enhancer context sensitivity delivering payloads to replace the *Sox2* LCR. **c, e.** *Sox2* expression analysis for payloads comprising combinations of *Sox2* 19, *Sox2* 20, *Sox2* 23, and *Sox2* 24. Each point represents *Sox2* expression from the engineered allele in an independent mESC clone relative to baseline payloads (dashed vertical lines). Bars indicate median. Expression was scaled between 0 (Δ*Sox2*) to 1 (WT LCR). Vertical dashed lines indicate median expression of baseline payloads. **d.** Average expression from (**c**). Additionally included previously published data for each DHS alone (gray) (Brosh et al. 2023).

**Fig. S9.**
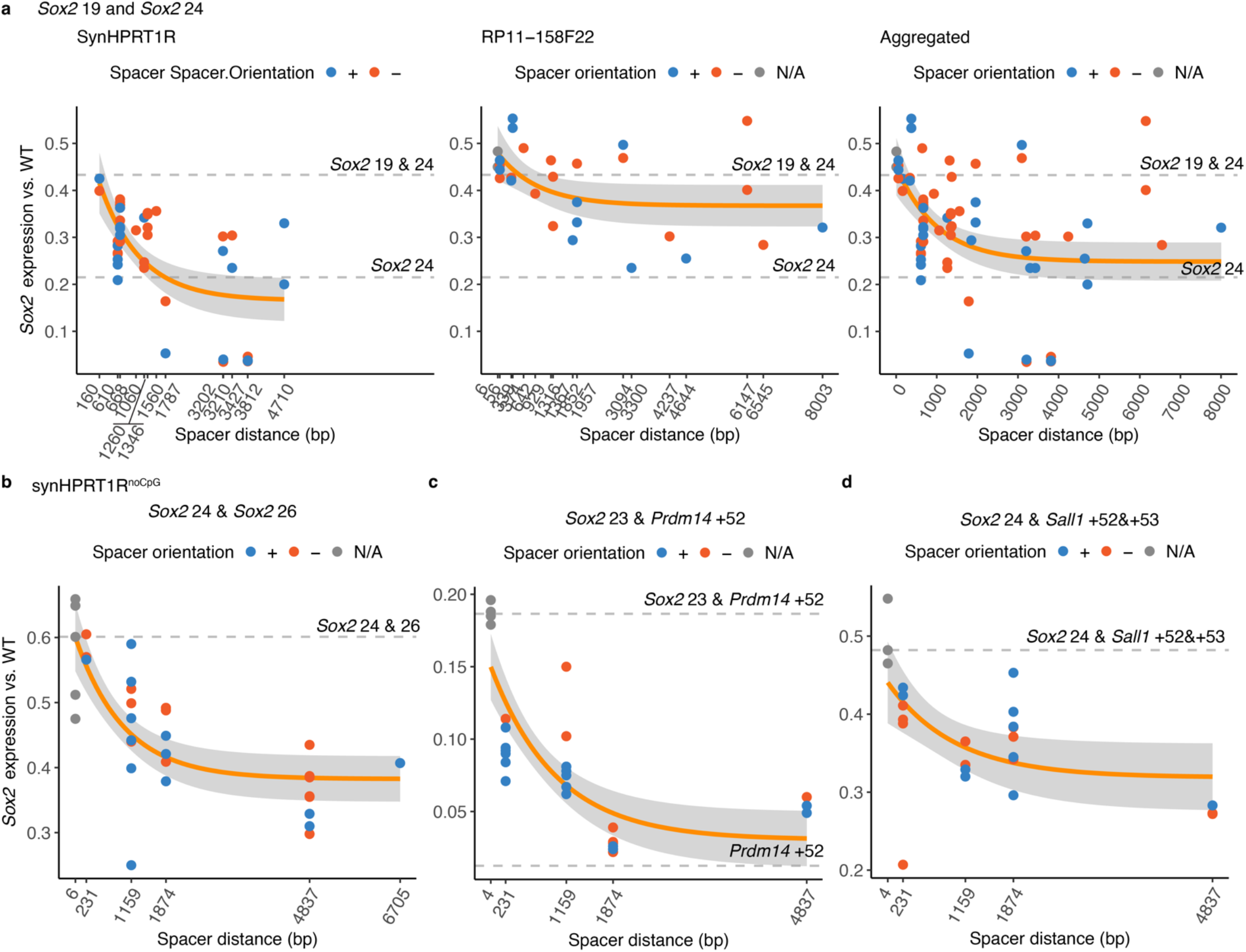
Extended analysis of distance-dependent synergy across DHS pairs. *Sox2* expression analysis using qRT-PCR in individual mESC clones including DHS pairs *Sox2* 19 and *Sox2* 24 (**a**), *Sox2* 24 and *Sox2* 26 (**c**), *Sox2* 23 and *Prdm14* +52 (**c**), and *Sox2* 24 and *Sall1* +52&+53 (**d**), each separated by spacers of varying lengths. Each point represents *Sox2* expression from the engineered allele in an independent mESC clone. Expression from the BL6 allele was normalized to the CAST allele and scaled between 0 (Δ*Sox2*) to 1 (WT LCR). Orange line indicates exponential fit with 95% confidence interval shaded in gray. Vertical dashed lines indicate median expression of baseline payloads. Color represents spacer orientation; N/A (grey) indicates that no spacer was cloned between the two DHSs and only the restriction enzyme sequence remains. **a**. *Sox2* 19 and *Sox2* 24 used spacers derived from SynHPRT1R and a human gene desert BAC, RP11-158F11. **b-d**. Other DHS pairs used spacer sequences from SynHPRT1R^noCpG^.

**Fig. S10.**
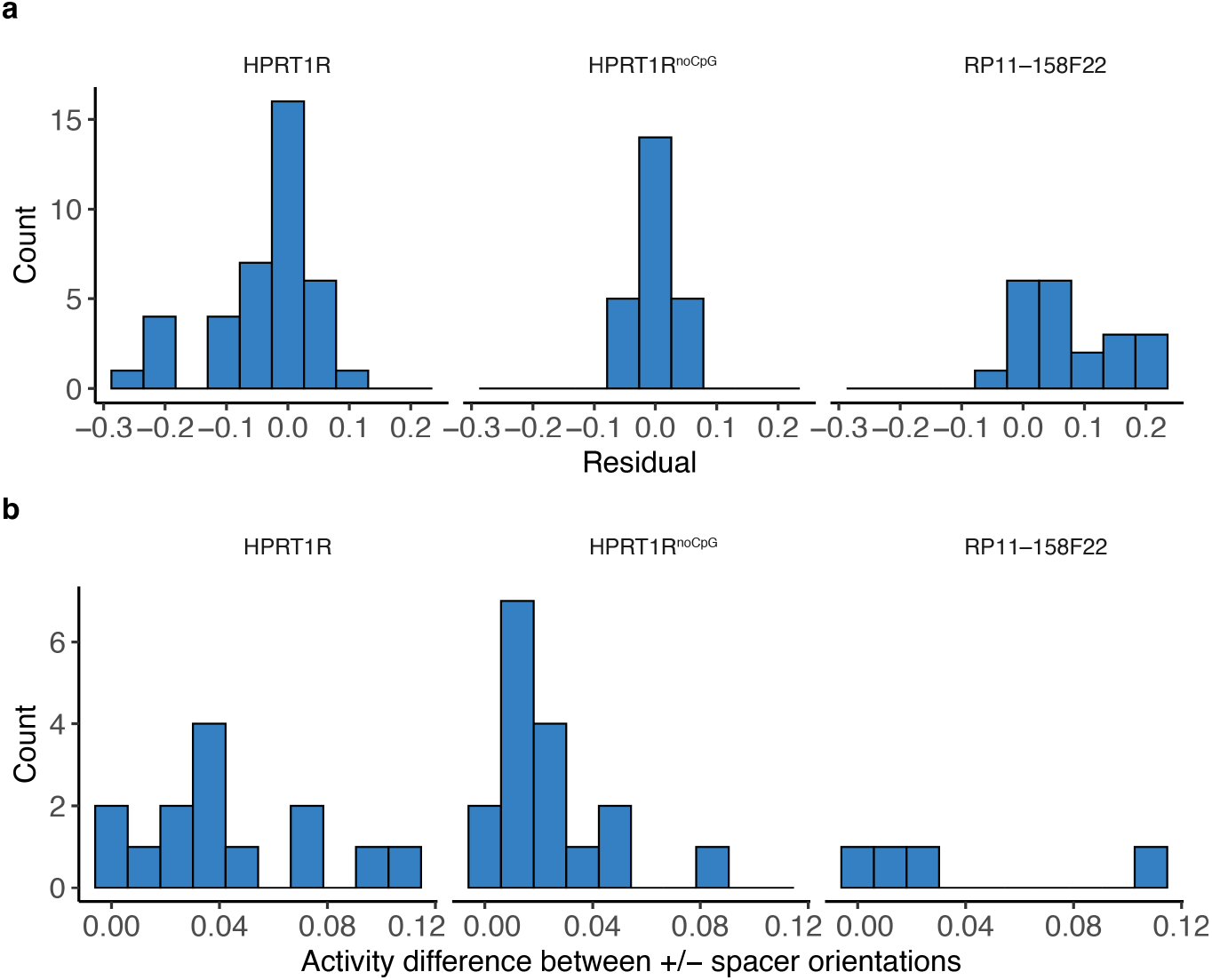
DHS spacer analysis residuals. Histograms of residuals from regressions in Fig. 5 and **Fig. S9** across all spacer fragments and orientations tested (**a**) and difference in activity between spacer orientations (**b**).

### SUPPLEMENTARY TABLES

**Table S1. Payload and assembly details.**

Strategies and reagents used for payload assembly, as well as sequencing data. Payload.ID represents a unique identifier for each payload. See **Table S5** and **Table S6** for payload labels used in Figures. Sample.ID, unique sequencing library identifier. Assembly strategy details are in **Methods**. See **Table S9** for cloning primer sequences and synthetic fragment sequences. UNS indicates the presence or absence of Unique Nucleotide Sequences flanking the payload. Intermediate assemblies that were not delivered are listed at the bottom. Comments identifies payloads that used slightly different DHS coordinates from a previous publication (Brosh et al. 2023) **(Fig. S4**).

**Table S2. Payload coordinates.**

Genomic coordinates (mm10) for payloads delivered in this study.

**Table S3. Payload composition for multiplexed delivery pools.**

Summary of different pools delivered in independent Big-IN experiments, including the specific payloads in each pool.

**Table S4. mESC clone summary.**

Summary table for mESC clones described in this paper. Spacer fragment assignment including size and orientation is listed for spacer-containing clones. Clone.ID represents a unique identifier for each mESC clone. See **Table S5** and **Table S6** for payload labels used in Figures. Paren-tal.Line represents the mESC clone the payloads were delivered to. Sample.ID, unique sequencing library identifier. Delivery.Pool.ID is “Individual” to denote arrayed deliveries or includes the pool ID for multiplexed deliveries (**Table S3**).

**Table S5. *Sox2* expression values and activity.**

*Sox2* expression in engineered mESC clones. The columns *Sox2* (CAST) and *Sox2* (BL6) list qRT-PCR ΔCt values relative to *Gapdh* for each *Sox2* allele. Fold.change indicates the BL6/CAST 2^ΔCt^ ratio. Activity indicates scaled fold change as detailed in **Methods**. Label, name as in Figure panels.

**Table S6. *Igf2*/*H19* expression values**

*Igf2*/*H19* expression in engineered mESC clones. The columns *H19* (Engineered) and *Igf2* (Engineered) list qRT-PCR ΔCt values relative to *Gapdh* for the engineered allele. Fold.change indicates the 2^-ΔCt^ value. Label, name as in Figure panels.

**Table S7. Capture sequencing libraries and quality control analysis.**

Summary table of DNA capture sequencing libraries. Sample.ID, a unique sequencing library ID; For each sample, the number (#) of sequenced, analyzed, and duplicate reads is provided. Bait.Set lists the capture baits employed. Genetic Modification details DNA elements integrated and their integration site in square brackets; transiently transfected plasmids (e.g. pSpCas9 or pCAG-Cre) are indicated by null. CoverageQC, IntegrationQC, KmerGenotypeQC, and Spac-erGenotypeQC indicate mESC clone quality control (QC) status as either PASS or FAIL; NA indicates not applicable.

**Table S8.** Trait-matched DNase-seq samples and SNPs overlapping DHSs.

| Trait Name | DNase Samples | Number of SNPs in DHSs |
| --- | --- | --- |
| Bilirubin (D) | Caco2-DS8235, HepG2-DS24838, hTH2-DS17597, fHeart_atrium-DS37460, HuH_7-ENCLB703KYC, HuH_75-ENCLB677WKD, fKidney_renal_cortex-DS20445 | 56 |
| Blood Pressure (Diastolic) | fAdrenal-DS19583, fKidney_renal_cortex-DS20445, fKidney_renal_pelvis-DS20448, fMuscle_lower_limb-DS18174, fHeart_atrium-DS37460, caput_mediale_musculus_gastrocnemius-DS37185 | 195 |
| Blood Pressure (Systolic) | fAdrenal-DS19583, fKidney_renal_cortex-DS20445, fKidney_renal_pelvis-DS20448, fMuscle_lower_limb-DS18174, fHeart_atrium-DS37460, caput_mediale_musculus_gastrocnemius-DS37185 | 221 |
| Calcium | Caco2-DS8235, HuH_7-ENCLB703KYC, HuH_75-ENCLB677WKD, CD20-DS17706, HepG2-DS24838 | 196 |
| Glucose | islet-SRR8729334, Caco2-DS8235, HuH_75-ENCLB677WKD, HuH_7-ENCLB703KYC, globus-DS23951 | 75 |
| Hypothyroidism | hTH17-DS11039, hTR-DS14702, hTH2-DS17597, hTH1-DS18018, CD3-DS17198, CD56-DS17189, CD8-DS17885, CD20-DS17706, CD4-DS17881, CD1c-DS31642, CD19-DS17186, CD14-DS17215, CD34-DS12734 | 152 |
| LDL Cholesterol | Caco2-DS8235, HepG2-DS24838, fLiver-DS23957, HuH_75-ENCLB677WKD, fMuscle_upperLimb-DS17661 | 197 |
| RBC Count | CD34_T18-DS25969, fLiver-DS23957, FLER-DS13348 | 392 |
| Triglycerides | Caco2-DS8235, HepG2-DS24838, fLiver-DS23957, HuH_75-ENCLB677WKD, fMuscle_upperLimb-DS17661 | 133 |
| Type 2 Diabetes | islet-SRR8729334, Caco2-DS8235, HuH_75-ENCLB677WKD, HuH_7-ENCLB703KYC, caput_mediale_musculus_gastrocnemius-DS37185, intestine_Sm-DS20770, adipocyte-DS26410 | 146 |

Traits analyzed in this study, associated matched DNase-seq samples, and number of unique fine-mapped trait-associated SNPs overlapping DHSs. Source data from (Forgetta et al. 2022).

**Table S9. Primers and synthetic DNA sequences.**

Oligonucleotides and synthetic DNA sequences used in this study, including cloning primers, synthetic fragments used for payload assembly, genotyping primers, and qRT-PCR primers.

**Table S10. mESC clone capture sequencing coverage quality control results.**

Sample.ID, unique sequencing library identifier. Average coverage depth is reported for custom references representing landing pad (LP), VectorBackbone, and pCAGiCre not expected to be present, as well as the flanking (the regions flanking the engineered site that were targeted by Capture-seq), payload and non-payload regions (the regions of the *Sox2* and *Igf2* loci included within and outside the payload, respectively). Coverage depth at payload and non-payload regions were normalized by coverage at flanking regions covered by same bait. CoverageQC represents overall quality assessment as PASS or FAIL (detailed in **Methods**).

**Table S11. mESC clone bamintersect results.**

Bamintersect integration site analysis for all clones. Each line describes a junction detected (detailed in **Methods**). Sample.ID, unique sequence library identifier. chrom (chromosome), chrom-Start, chromEnd, Width, NearestGene, and Strand are shown; bam1 and bam2 correspond to reference and custom reference, respectively. NearestGene indicates the nearest GENCODE v23 gene and its distance in bp (negative values indicate an upstream element) or other annotated element. ReadsPer10M, number of read pairs supporting the junction normalized to 10 million sequenced reads. Genomes indicates the reference and custom genomes compared. Jx lists the junction classification as LeftJx, RightJx, or endogenous (detailed in **Methods**).

**Table S12. mESC clone spacer bamintersect genotyping results.**

Bamintersect spacer genotyping analysis. Each line describes an internal junction detected between the payload DHS and synthetic spacer (detailed in **Methods**). Sample.ID, unique sequence library identifier; chromStart (start coordinate), chromEnd (end coordinate), Width, and Strand for bam1 and bam2, both corresponding to the payload custom reference. ReadsPer10M, number of read pairs supporting the junction normalized to 10 million; Jx, junction classification as LeftJx or RightJx (detailed in **Methods**); and JxStrand, indicates the spacer strand orientation.

**Table S13. mESC clone k-mer genotyping results.**

Pooled delivered (n=274), individually delivered (n=10), and small spacer (n=6) mESC clones k-mer genotyping results. Sample.ID, unique sequencing library identifier. KmerGenotype, genotyped Payload.ID. KmerSupport, read counts for k-mer evidence supporting inferred genotype. KmerConfounder, read counts for k-mer evidence inconsistent with inferred genotype.

**Table S14. K-mer sequences used for genotyping analysis.**

Collection of k-mers used in the k-mer genotyping analysis and their sequences.

**Table S15. Payload to k-mer assignments for mESC genotyping analysis.**

Payload to k-mer associations used for mESC k-mer genotyping, where each payload is identified by a unique set of k-mers (defined in **Table S14**).

### SUPPLEMENTARY DATA

#### Data S1. Payload sequences

Sequences for payloads described in this paper, provided in fasta format. Sequences include the region from LoxM to LoxP. See **Table S5** and **Table S6** for payload labels used in Figures.

#### Data S2. Raw qRT-PCR Cp values

Raw qRT-PCR Cp values for all mESC clones and assays. Source, the date when the assay was performed; Cp, threshold cycle.

## REFERENCES

1. Avsec Ž, Weilert M, Shrikumar A, Krueger S, Alexandari A, Dalal K, Fropf R, McAnany C, Gagneur J, Kundaje A, et al. 2021. Base-resolution models of transcription-factor binding reveal soft motif syntax. Nat Genet 53: 354–366.

2. Banerji J, Rusconi S, Schaffner W. 1981. Expression of a beta-globin gene is enhanced by remote SV40 DNA sequences. Cell 27: 299–308.

3. Batut PJ, Bing XY, Sisco Z, Raimundo J, Levo M, Levine MS. 2022. Genome organization controls transcriptional dynamics during development. Science 375: 566–570.

4. Bell AC, Felsenfeld G. 2000. Methylation of a CTCF-dependent boundary controls imprinted expression of the Igf2 gene. Nature 405: 482–485.

5. Benner C, Spencer CCA, Havulinna AS, Salomaa V, Ripatti S, Pirinen M. 2016. FINEMAP: efficient variable selection using summary data from genome-wide association studies. Bioinformatics 32: 1493–1501.

6. Blayney JW, Francis H, Rampasekova A, Camellato B, Mitchell L, Stolper R, Cornell L, Babbs C, Boeke JD, Higgs DR, et al. 2023. Super-enhancers include classical enhancers and facilitators to fully activate gene expression. Cell 186: 5826–5839.e18.

7. Blinka S, Reimer MH, Pulakanti K, Rao S. 2016. Super-Enhancers at the Nanog Locus Differentially Regulate Neighboring Pluripotency-Associated Genes. Cell Rep 17: 19–28.

8. Bothma JP, Garcia HG, Ng S, Perry MW, Gregor T, Levine M. 2015. Enhancer additivity and non-additivity are determined by enhancer strength in the Drosophila embryo. eLife 4.

9. Bower G, Hollingsworth EW, Jacinto SH, Alcantara JA, Clock B, Cao K, Liu M, Dziulko A, Alcaina-Caro A, Xu Q, et al. 2025. Range extender mediates long-distance enhancer activity. Nature 643: 830–838.

10. Brosh R, Coelho C, Ribeiro-Dos-Santos AM, Ellis G, Hogan MS, Ashe HJ, Somogyi N, Ordoñez R, Luther RD, Huang E, et al. 2023. Synthetic regulatory genomics uncovers enhancer context dependence at the Sox2 locus. Mol Cell 83: 1140–1152.e7.

11. Brosh R, Laurent JM, Ordoñez R, Huang E, Hogan MS, Hitchcock AM, Mitchell LA, Pinglay S, Cadley JA, Luther RD, et al. 2021. A versatile platform for locus-scale genome rewriting and verification. Proc Natl Acad Sci U S A 118: e2023952118.

12. Camellato BR, Brosh R, Ashe HJ, Maurano MT, Boeke JD. 2024. Synthetic reversed sequences reveal default genomic states. Nature 628: 373–380.

13. Catarino RR, Stark A. 2018. Assessing sufficiency and necessity of enhancer activities for gene expression and the mechanisms of transcription activation. Genes Dev 32: 202–223.

14. Chen L-F, Long HK, Park M, Swigut T, Boettiger AN, Wysocka J. 2023. Structural elements promote architectural stripe formation and facilitate ultra-long-range gene regulation at a human disease locus. Mol Cell 83: 1446–1461.e6.

15. Choi J, Lysakovskaia K, Stik G, Demel C, Söding J, Tian TV, Graf T, Cramer P. 2021. Evidence for additive and synergistic action of mammalian enhancers during cell fate determination. Elife 10: e65381.

16. Corradin O, Saiakhova A, Akhtar-Zaidi B, Myeroff L, Willis J, Cowper-Sal lari R, Lupien M, Mar-kowitz S, Scacheri PC. 2014. Combinatorial effects of multiple enhancer variants in linkage disequilibrium dictate levels of gene expression to confer susceptibility to common traits. Genome Research 24: 1–13.

17. ENCODE Project Consortium. 2012. An integrated encyclopedia of DNA elements in the human genome. Nature 489: 57–74.

18. Etchegaray J-P, Zhong L, Li C, Henriques T, Ablondi E, Nakadai T, Van Rechem C, Ferrer C, Ross KN, Choi J-E, et al. 2019. The Histone Deacetylase SIRT6 Restrains Transcription Elongation via Promoter-Proximal Pausing. Mol Cell 75: 683–699.e7.

19. Forgetta V, Jiang L, Vulpescu NA, Hogan MS, Chen S, Morris JA, Grinek S, Benner C, Jang D-K, Hoang Q, et al. 2022. An effector index to predict target genes at GWAS loci. Hum Genet 141: 1431–1447.

20. Grant CE, Bailey TL, Noble WS. 2011. FIMO: scanning for occurrences of a given motif. Bioinformatics 27: 1017–1018.

21. Halow JM, Byron R, Hogan MS, Ordoñez R, Groudine M, Bender MA, Stamatoyannopoulos JA, Maurano MT. 2021. Tissue context determines the penetrance of regulatory DNA variation. Nat Commun 12: 2850.

22. Hark AT, Schoenherr CJ, Katz DJ, Ingram RS, Levorse JM, Tilghman SM. 2000. CTCF mediates methylation-sensitive enhancer-blocking activity at the H19/Igf2 locus. Nature 405: 486–489.

23. Hnisz D, Abraham BJ, Lee TI, Lau A, Saint-André V, Sigova AA, Hoke HA, Young RA. 2013. Super-enhancers in the control of cell identity and disease. Cell 155: 934–947.

24. Hnisz D, Schuijers J, Lin CY, Weintraub AS, Abraham BJ, Lee TI, Bradner JE, Young RA. 2015. Convergence of developmental and oncogenic signaling pathways at transcriptional su-per-enhancers. Mol Cell 58: 362–370.

25. Jensen CL, Chen L-F, Swigut T, Crocker OJ, Yao D, Bassik MC, Ferrell JE, Boettiger AN, Wysocka J. 2025. Long-range regulation of transcription scales with genomic distance in a gene-specific manner. Mol Cell 85: 347–361.e7.

26. Jolma A, Yan J, Whitington T, Toivonen J, Nitta KR, Rastas P, Morgunova E, Enge M, Taipale M, Wei G, et al. 2013. DNA-Binding Specificities of Human Transcription Factors. Cell 152: 327–339.

27. Karr JP, Ferrie JJ, Tjian R, Darzacq X. 2022. The transcription factor activity gradient (TAG) model: contemplating a contact-independent mechanism for enhancer-promoter communication. Genes Dev 36: 7–16.

28. Kassouf MT, Francis HS, Gosden M, Suciu MC, Downes DJ, Harrold C, Larke M, Oudelaar M, Cornell L, Blayney J, et al. 2025. The α-globin super-enhancer acts in an orientation-dependent manner. Nat Commun 16: 1033.

29. Koeppel J, Murat P, Girling G, Peets EM, Gouley M, Rebernig V, Maheshwari A, Hepkema J, Weller J, Jebaraj JHJ, et al. 2025. Resolution of a human super-enhancer by targeted genome randomisation. bioRxiv 2025.01.14.632548.

30. Li B, Dewey CN. 2011. RSEM: accurate transcript quantification from RNA-Seq data with or without a reference genome. BMC Bioinformatics 12: 323.

31. Li Q, Peterson KR, Fang X, Stamatoyannopoulos G. 2002. Locus control regions. Blood 100: 3077–3086.

32. Li Y, Rivera CM, Ishii H, Jin F, Selvaraj S, Lee AY, Dixon JR, Ren B. 2014. CRISPR reveals a distal super-enhancer required for Sox2 expression in mouse embryonic stem cells. PLoS One 9: e114485.

33. Lin X, Liu Y, Liu S, Zhu X, Wu L, Zhu Y, Zhao D, Xu X, Chemparathy A, Wang H, et al. 2022. Nested epistasis enhancer networks for robust genome regulation. Science 377: 1077–1085.

34. Long E, Williams J, Zhang H, Choi J. 2025. An evolving understanding of multiple causal variants underlying genetic association signals. Am J Hum Genet 112: 741–750.

35. Love MI, Huber W, Anders S. 2014. Moderated estimation of fold change and dispersion for RNA-seq data with DESeq2. Genome Biol 15: 550.

36. Martinez-Ara M, Comoglio F, van Arensbergen J, van Steensel B. 2022. Systematic analysis of intrinsic enhancer-promoter compatibility in the mouse genome. Mol Cell 82: 2519–2531.e6.

37. Maurano MT. 2026. Synthetic Regulatory Genomics. Annu Rev Genomics Hum Genet.

38. Maurano MT, Haugen E, Sandstrom R, Vierstra J, Shafer A, Kaul R, Stamatoyannopoulos JA. 2015. Large-scale identification of sequence variants influencing human transcription factor occupancy in vivo. Nature Genetics 47: 1393–1401.

39. Maurano MT, Humbert R, Rynes E, Thurman RE, Haugen E, Wang H, Reynolds AP, Sandstrom R, Qu H, Brody J, et al. 2012. Systematic localization of common disease-associated variation in regulatory DNA. Science 337: 1190–1195.

40. Meuleman W, Muratov A, Rynes E, Halow J, Lee K, Bates D, Diegel M, Dunn D, Neri F, Teodo-siadis A, et al. 2020. Index and biological spectrum of human DNase I hypersensitive sites. Nature 584: 244–251.

41. Mirny LA. 2010. Nucleosome-mediated cooperativity between transcription factors. Proc Natl Acad Sci U S A 107: 22534–22539.

42. Moorthy SD, Davidson S, Shchuka VM, Singh G, Malek-Gilani N, Langroudi L, Martchenko A, So V, Macpherson NN, Mitchell JA. 2017. Enhancers and super-enhancers have an equivalent regulatory role in embryonic stem cells through regulation of single or multiple genes. Genome Res 27: 246–258.

43. Neph S, Kuehn MS, Reynolds AP, Haugen E, Thurman RE, Johnson AK, Rynes E, Maurano MT, Vierstra J, Thomas S, et al. 2012. BEDOPS: high-performance genomic feature operations. Bioinformatics 28: 1919–1920.

44. Ordoñez R, Zhang W, Ellis G, Zhu Y, Ashe HJ, Ribeiro-Dos-Santos AM, Brosh R, Huang E, Hogan MS, Boeke JD, et al. 2024. Genomic context sensitizes regulatory elements to genetic disruption. Mol Cell 84: 1842–1854.e7.

45. Parker SCJ, Stitzel ML, Taylor DL, Orozco JM, Erdos MR, Akiyama JA, van Bueren KL, Chines PS, Narisu N, NISC Comparative Sequencing Program, et al. 2013. Chromatin stretch enhancer states drive cell-specific gene regulation and harbor human disease risk variants. Proceedings of the National Academy of Sciences of the United States of America 110: 17921–17926.

46. Peng T, Zhai Y, Atlasi Y, Ter Huurne M, Marks H, Stunnenberg HG, Megchelenbrink W. 2020. STARR-seq identifies active, chromatin-masked, and dormant enhancers in pluripotent mouse embryonic stem cells. Genome Biol 21: 243.

47. Ramírez F, Ryan DP, Grüning B, Bhardwaj V, Kilpert F, Richter AS, Heyne S, Dündar F, Manke T. 2016. deepTools2: a next generation web server for deep-sequencing data analysis. Nucleic Acids Res 44: W160–165.

48. Ribeiro-Dos-Santos AM, Hogan MS, Luther RD, Brosh R, Maurano MT. 2022. Genomic context sensitivity of insulator function. Genome Res 32: 425–436.

49. Schaffner W. 2015. Enhancers, enhancers - from their discovery to today’s universe of transcription enhancers. Biol Chem 396: 311–327.

50. Singh G, Mullany S, Moorthy SD, Zhang R, Mehdi T, Tian R, Duncan AG, Moses AM, Mitchell JA. 2021. A flexible repertoire of transcription factor binding sites and a diversity threshold determines enhancer activity in embryonic stem cells. Genome Res 31: 564–575.

51. Small S, Arnosti DN, Levine M. 1993. Spacing ensures autonomous expression of different stripe enhancers in the even-skipped promoter. Development 119: 762–772.

52. Sobreira DR, Joslin AC, Zhang Q, Williamson I, Hansen GT, Farris KM, Sakabe NJ, Sinnott-Armstrong N, Bozek G, Jensen-Cody SO, et al. 2021. Extensive pleiotropism and allelic heterogeneity mediate metabolic effects of IRX3 and IRX5. Science 372: 1085–1091.

53. Stergachis AB, Debo BM, Haugen E, Churchman LS, Stamatoyannopoulos JA. 2020. Single-molecule regulatory architectures captured by chromatin fiber sequencing. Science 368: 1449–1454.

54. Swanson EG, Mao Y, Mallory BJ, Vollger MR, Bohaczuk SC, Oliveira CB, Lyon DB, Ranchalis J, Parmalee NL, Cohen BA, et al. 2025. Mapping single-cell diploid chromatin fiber architectures using DAF-seq. Nat Biotechnol.

55. The ENCODE Project Consortium, Moore JE, Purcaro MJ, Pratt HE, Epstein CB, Shoresh N, Adrian J, Kawli T, Davis CA, Dobin A, et al. 2020. Expanded encyclopaedias of DNA elements in the human and mouse genomes. Nature 583: 699–710.

56. Thomas HF, Feng S, Haslhofer F, Huber M, García Gallardo M, Loubiere V, Vanina D, Pitasi M, Stark A, Buecker C. 2025. Enhancer cooperativity can compensate for loss of activity over large genomic distances. Mol Cell 85: 362–375.e9.

57. Thomas HF, Kotova E, Jayaram S, Pilz A, Romeike M, Lackner A, Penz T, Bock C, Leeb M, Halbritter F, et al. 2021. Temporal dissection of an enhancer cluster reveals distinct temporal and functional contributions of individual elements. Mol Cell 81: 969–982.e13.

58. Thurman RE, Rynes E, Humbert R, Vierstra J, Maurano MT, Haugen E, Sheffield NC, Stergachis AB, Wang H, Vernot B, et al. 2012. The accessible chromatin landscape of the human genome. Nature 489: 75–82.

59. Vierstra J, Rynes E, Sandstrom R, Zhang M, Canfield T, Hansen RS, Stehling-Sun S, Sabo PJ, Byron R, Humbert R, et al. 2014. Mouse regulatory DNA landscapes reveal global principles of cis-regulatory evolution. Science 346: 1007–1012.

60. Vos ESM, Valdes-Quezada C, Huang Y, Allahyar A, Verstegen MJAM, Felder A-K, van der Vegt F, Uijttewaal ECH, Krijger PHL, de Laat W. 2021. Interplay between CTCF boundaries and a super enhancer controls cohesin extrusion trajectories and gene expression. Mol Cell 81: 3082–3095.e6.

61. Whyte WA, Orlando DA, Hnisz D, Abraham BJ, Lin CY, Kagey MH, Rahl PB, Lee TI, Young RA. 2013. Master transcription factors and mediator establish super-enhancers at key cell identity genes. Cell 153: 307–319.

62. Yue F, Cheng Y, Breschi A, Vierstra J, Wu W, Ryba T, Sandstrom R, Ma Z, Davis C, Pope BD, et al. 2014. A comparative encyclopedia of DNA elements in the mouse genome. Nature 515: 355–364.

63. Zhou HY, Katsman Y, Dhaliwal NK, Davidson S, Macpherson NN, Sakthidevi M, Collura F, Mitchell JA. 2014. A Sox2 distal enhancer cluster regulates embryonic stem cell differentiation potential. Genes & Development 28: 2699–2711.

64. Zhou Z, Cheng Y, Lieberth J, Zhang J, Jin Y, Li A, Yu H, Ozer A, Lis JT. 2026. Robust regulatory interplay of enhancers, facilitators, and promoters in a native chromatin context. Cell S0092-8674(26)00753–1.

65. Zuin J, Roth G, Zhan Y, Cramard J, Redolfi J, Piskadlo E, Mach P, Kryzhanovska M, Tihanyi G, Kohler H, et al. 2022. Nonlinear control of transcription through enhancer-promoter interactions. Nature 604: 571–577.

